# The architecture of metabolism maximizes biosynthetic diversity in the largest class of fungi

**DOI:** 10.1101/2020.01.31.928846

**Authors:** Emile Gluck-Thaler, Sajeet Haridas, Manfred Binder, Igor V. Grigoriev, Pedro W. Crous, Joseph W. Spatafora, Kathryn Bushley, Jason C. Slot

## Abstract

**Background:** Ecological diversity in fungi is largely defined by metabolic traits, including the ability to produce secondary or “specialized” metabolites (SMs) that mediate interactions with other organisms. Fungal SM pathways are frequently encoded in biosynthetic gene clusters (BGCs), which facilitate the identification and characterization of metabolic pathways. Variation in BGC composition reflects the diversity of their SM products. Recent studies have documented surprising diversity of BGC repertoires among isolates of the same fungal species, yet little is known about how this population-level variation is inherited across macroevolutionary timescales.

**Results:** Here, we applied a novel linkage-based algorithm to reveal previously unexplored dimensions of diversity in BGC composition, distribution, and repertoire across 101 species of Dothideomycetes, which are considered to be the most phylogenetically diverse class of fungi and are known to produce many SMs. We predicted both complementary and overlapping sets of clustered genes compared with existing methods and identified novel gene pairs that associate with known secondary metabolite genes. We found that variation in BGC repertoires is due to non-overlapping BGC combinations and that several BGCs have biased ecological distributions, consistent with niche-specific selection. We observed that total BGC diversity scales linearly with increasing repertoire size, suggesting that secondary metabolites have little structural redundancy in individual fungi.

**Conclusions:** We project that there is substantial unsampled BGC diversity across specific families of Dothideomycetes, which will provide a roadmap for future sampling efforts. Our approach and findings lend new insight into how BGC diversity is generated and maintained across an entire fungal taxonomic class.

## Background

Plants, bacteria and fungi produce the majority of the earth’s biochemical diversity. These organisms produce a remarkable variety of secondary/specialized metabolites (SMs) that can mediate ecological functions, including defense, resource acquisition, and mutualism. Standing SM diversity is often high at the population level, which may affect the rates of adaptation over microevolutionary timescales. For example, high intraspecific quantitative and qualitative chemotype diversity in plants can enable rapid adaptation to local biotic factors (Agrawal, Hastings et al. 2012; Züst, Heichinger et al. 2012; Glassmire, Jeffrey et al. 2016). However, the fate of population-level chemodiversity across longer timescales is not well explored in plants or other lineages. We therefore sought to identify how metabolic variation is distributed across macroevolutionary timescales by profiling chemodiversity across a well-sampled taxonomic class.

The Dothideomycetes, which originated between 247 and 459 million years ago (Beimforde, Feldberg et al. 2014), comprise the largest and arguably most phylogenetically diverse class of fungi. Currently, 19,000 species are recognized in 32 orders containing more than 1,300 genera (Zhang, Crous et al. 2011). Dothideomycetes are divided into two major subclasses, the Pleosporomycetidae (order Pleosporales) and Dothideomycetidae (orders Dothideales, Capnodiales, and Myriangiales), which correspond to the presence or absence, respectively, of pseudoparaphyses during development of the asci (Schoch, Crous et al. 2009). Several other orders await definitive placement.

Dothideomycetes also display a large diversity of fungal lifestyles and ecologies. The majority of Dothideomycetes are terrestrial and associate with phototrophic hosts as either pathogens, saprobes, endophytes (Schoch, Crous et al. 2009), lichens (Nelsen, Lucking et al. 2011), or ectomycorrhizal symbionts (Spatafora, Owensby et al. 2012). Six orders contain plant pathogens capable of infecting nearly every known crop species. The Pleosporales and Capnodiales, in particular, are dominated by asexual plant pathogens that cause significant economic losses and have been well sampled in previous genome sequencing efforts (Goodwin, Ben M’Barek et al. 2011; Ohm, Feau et al. 2012; Oliver, Friesen et al. 2012; Condon, Leng et al. 2013; Manning, Pandelova et al. 2013). A single order (Jahnulales) contains aquatic, primarily freshwater, species (Suetrong, Boonyuen et al. 2011). Other ecologies include human and animal pathogens, including some taxa that can elicit allergies and asthma (Crameri, Garbani et al. 2014), and rock-inhabiting fungi (Ruibal, Gueidan et al. 2009).

This broad range of lifestyles is accompanied by extensive diversity of SMs, for which very few have known ecological roles. The Dothideomycetes, and several other ascomycete classes (Eurotiomycetes, Sordariomycetes, and Leotiomycetes), produce the greatest number and diversity of SMs across the fungal kingdom (Spatafora and Bushley 2015; Akimitsu et al. 2014). Economically important plant pathogens in the Pleosporales (*Alternaria, Bipolaris, Exserohilum, Leptosphaeria, Pyrenophora,* and *Stagonospora*), in particular, are known to produce host-selective toxins that confer the ability to cause disease in specific plant hosts (Walton and Panaccione 1993; Walton 1996; Wolpert, Dunkle et al. 2002; Ciuffetti, Manning et al. 2010, Pandelova, Figueroa et al. 2012; Akimitsu, Tsuge et al. 2014). Other toxins first identified in Pleosporales have roles in virulence, but are not pathogenicity determinants, including the PKS derived compounds depudecin (Wight, Kim et al. 2009) and solanapyrone (Kaur 1995).

Dothideomycetes are also known to produce bioactive metabolites shared with more distantly related fungal classes. Sirodesmin, a virulence factor produced by *Leptosphaeria maculans,* for example, belongs to the same class of epipolythiodioxopiperazine (ETP) toxins as gliotoxin, an immunosuppressant produced by the eurotiomycete human pathogen *Aspergillus fumigatus* (Gardiner, Waring et al. 2005; Patron, Waller et al. 2007). Dothistromin, a polyketide metabolite produced by the pine pathogen *Dothistroma septosporum* shares ancestry with aflatoxin (Bradshaw, Slot et al. 2013), a mycotoxin produced by *Aspergillus* species that poses serious human health and environmental risks worldwide (Horn 2003; Wang and Tang 2004).

A majority of bioactive metabolites in Dothideomycetes are small-molecule SMs that, like those of other fungi, are frequently the products of biosynthetic gene clusters (BGCs) composed of enzymes, transporters, and regulators that contribute to a common SM pathway. Most of these BGCs are defined by four main classes of SM core signature enzymes: 1) nonribosomal peptide synthetases (NRPS), 2) polyketide synthetases (PKS), 3) terpene synthases (TS), and 4) dimethylallyl tryptophan synthases (DMAT) (Hoffmeister & Keller 2007). Fungal gene clusters are hotspots for genome evolution through gene duplication, loss, and horizontal transfer, which recombine pathways and generate diversity (Wisecaver, Slot et al. 2014). Additionally, recent studies have shown that gene clusters may evolve through recombination or shuffling of modular subunits of syntenic genes (Lind, Wisecaver et al. 2017; Gluck-Thaler et al 2018. Changes in BGC gene content often result in structural changes to the SM product(s), and therefore BGCs can be used to monitor the evolution of chemodiversity (Lind, Wisecaver et al. 2017; Proctor, McCormick et al. 2018). The most widely used methods for detecting BGCs rely on models of gene cluster composition based on putative functions in SM biosynthesis informed by a phylogenetically limited set of taxa, but gene function agnostic methods are being developed (Slot and Gluck-Thaler 2019).

Here, we systematically assessed BGC richness and compositional diversity in the genomes of 101 Dothideomycetes species, most recently sequenced (Haridas et al. *in press*). Using a newly benchmarked algorithm that identifies clustered genes of interest through the frequency of their co-occurrence with and around signature biosynthetic genes, we identified 3399 putative BGCs, grouped into 719 unique cluster types, including 5 varieties of candidate DHN melanin clusters. The conservation of specific gene pairs across BGC types suggests that precise functional interactions contribute to the modular evolution of these loci. Numerous BGCs have either over- or under-dispersed phylogenetic distributions, suggesting pathways have been differentially impacted by selection. In comparisons across species, BGC repertoire diversity increases linearly with repertoire size, reflecting a mode of metabolic evolution in these fungi that is likely distinct from that of plants. We found little overlap in cluster repertoires among genomes from different genera, and project that a wealth of unique BGCs remain to be discovered within this fungal lineage.

## Results

### Dothideomycetes contain hundreds of distinct types of BGCs, a small fraction of which are characterized

Using a novel cluster detection approach based on shared syntenic relationships among genes (CO-OCCUR, see Methods, Figure 1, Figure SA), we identified 332 gene homolog groups (homolog groups) of interest (Table SA, Table SB) whose members were organized into 3399 candidate BGCs of at least two genes (Table SC) in 101 Dothideomycete genomes (Table SD), representing an average of 33.7 BGCs per genome (SD= 15.4, Figure SG). We grouped BGCs into 719 unique cluster types based on a minimum gene content similarity of 90%; 422 cluster types are part of homologous cluster groups (**cluster group**s) found in 2 or more genomes, and 297 are orphan clusters found in only one genome (Table SE). Of these, 345 cluster types (166 cluster groups and 179 orphan clusters) had 5 or more genes per BGC (Figure 2), and 459 cluster types (239 cluster groups and 220 orphan clusters) had 4 or more genes per BGC (Figure SC). Only 9 of the 459 cluster types with greater than 4 genes were ever found more than once in any given genome (Table SE). According to standard practice, we classified cluster types based on the presence of biosynthetic signature genes: dimethylallyl tryptophan synthase (DMAT), polyketide synthase (PKS), PKS-like, nonribosomal peptide-synthetase (NRPS), NRPS-like, hybrid (HYBRID), and terpene cyclase (TC). We found that among all cluster types with greater than 4 genes, 186 contained only PKS and 29 contained only NRPS signature genes. Similarly, we detected 4 DMAT, 38 PKS-like, 16 NRPS-like, 3 HYBRID, and 3 TC-only cluster types. 127 cluster types contained more than 1 type of signature gene, and 53 cluster types contained no signature gene at all but still consisted of genes found in significant co-occurrences. By searching the MIBiG database for highly similar hits (≥70% amino acid identity) to the signature biosynthetic genes in CO-OCCUR BGCs, we were able to confidently annotate 158 of the BGCs recovered by CO-OCCUR with 32 unique MIBiG entries, corresponding to 22 unique metabolites (Table SF). BGC annotations based instead on content overlap with characterized MIBiG clusters can be found in Table SG (minimum cluster size=3 genes; minimum percentage of genes with similarity=70%).

**Figure 1.**
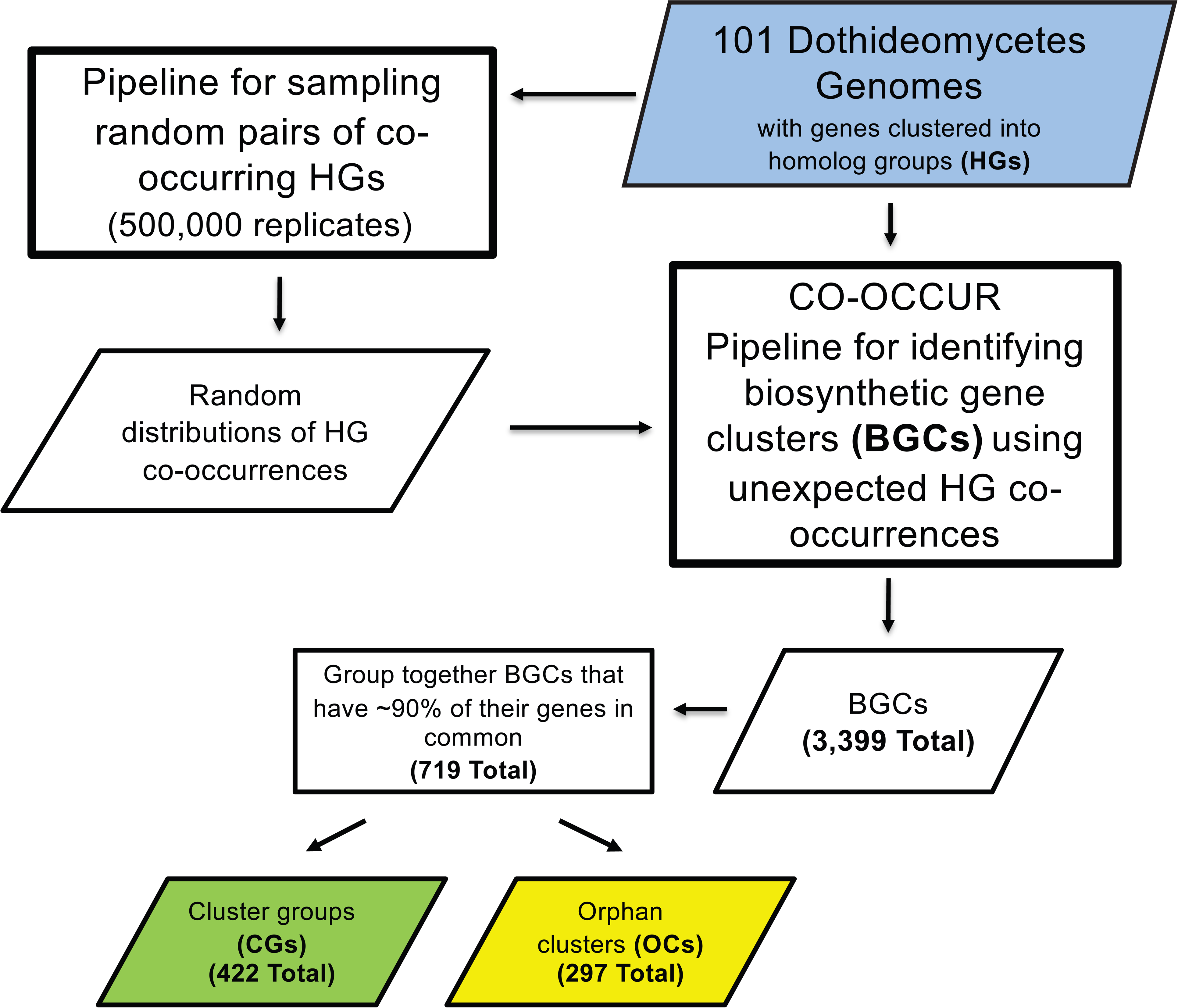
CO-OCCUR pipeline. The pipeline used genome annotations from 101 Dothideomycetes, and previously computed homolog groups (HGs) consisting of both orthologs and paralogs (Haridas et al. *in press*). Biosynthetic gene clusters (BGCs) were inferred by determining unexpectedly distributed shared HG pairs, determined according to a null-distribution of randomly sampled gene pairs in the same genomes, and then a search for all clusters containing the HG pairs. The resulting BGCs were then either consolidated into cluster groups (CGs) that share ∼90% of gene content or labeled orphan cluster (OCs) if found only in a single taxon. A detailed pipeline is presented in Figure SA.

**Figure 2.**
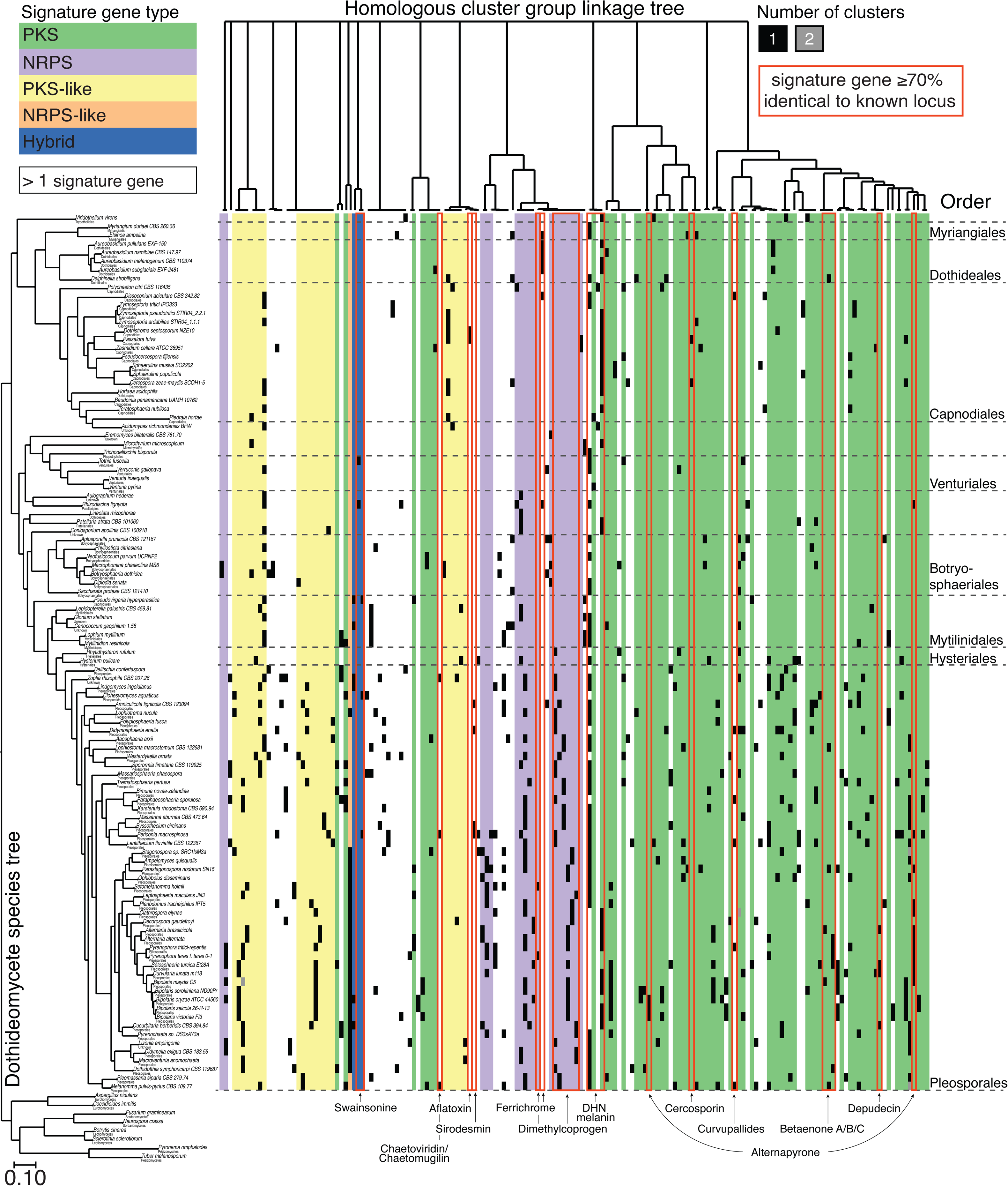
Diversity of the largest detected secondary metabolite gene clusters across 101 Dothideomycetes. A maximum likelihood phylogenomic tree of 101 Dothideomycete species (Haridas et al. *in press*) corresponds to rows in a heatmap (right) that depicts the number of secondary metabolite clusters found in each genome, delimited by order (dotted line). Each cluster is assigned to a homologous cluster group (cluster group; column) defined by at least 90% similarity at the composition level. Only cluster groups with ≥ 5 unique homolog groups per cluster are shown. A complete linkage tree (top) depicts relationships among cluster groups, where distance is proportional to the Raup-Crick dissimilarity in cluster group composition. Cluster groups are colored according to their core signature biosynthetic genes, and cluster groups with greater than 1 signature gene are left uncolored. Cluster groups with signature genes ≥70% identical to characterized BGC signature genes in MIBiG are indicated by a labeled red box.

Of the 158 BGCs with hits to the MIBiG database, some encoded non-host selective phytotoxins or other compounds with known roles in virulence to plants. Two PKS BGC’s encoding the non-host-specific phytotoxin and DNA polymerase inhibitor, solanapyrone (Mizushina, Kamisuki et al. 2002, Kasahara, Miyamoto et al. 2010), and the related alternapyrone (Fujii, Yoshida et al. 2005), first identified in *Alternaria solani* (Kasahara et al. 2010; Mizushina et al. 2002; Fujii et al. 2005), were found across taxa primarily in the order Pleosporales, especially in the closely related Pleosporaceae, Leptosphaeriaceae, and Phaeosphaeriaceae families (Figure 2, Table SF, SG). Several BGCs mapping to the MIBiG cluster for the extracellular siderophore dimethylcoprogen, which plays a role in virulence in the corn pathogen *Cochliobolus heterostrophus* (Dothideomycetes) and *Fusarium graminearum* (Sordariomycetes), was also found in most taxa in Pleosporales (Oide, Moeder et al. 2006). In contrast, a BGC mapping to the NRPS phytotoxin sirodesmin, produced by *Leptosphaeria maculans* (Gardiner, Cozijnsen et al. 2004), and depudecin, a histone deacetylase (HDAC) synthesized by a PKS BGC first identified in *A. brassicicola* (Wight et al. 2009), were found discontinuously distributed in only a few unrelated species within Pleosporales (Figure 2, Table SF, SG). Aside from sirodesmin, only one other BGC had hits in MIBiG to a BGC producing a host-selective toxin. A BGC mapping to T-toxin, a polyketide toxin produced by race T (C4) of *C. heterostrophus* (only Race O (C5) included in this study) that was responsible for the devastating Southern Corn Leaf Blight (Daly 1982; Turgeon and Baker 2007), was detected by CO-OCCUR in only two additional taxa, *Ampelomyces quisqualis* and *L. maculans* (Table SG).

Other BGCs matched MIBiG clusters from other ascomycete classes (Eurotiomycetes, Sordariomycetes), some of which have been previously detected in Dothideomycetes while others were unexpected. The aflatoxin-like dothistromin clusters, which are fragmented into six mini-clusters in *Dothistroma septorum* (Bradshaw, Slot et al. 2013), predictably mapped to clusters detected by CO-OCCUR in *D. septorum* <u>and the closely related species</u> *Passalora fulva* (Capnodiales) Some unexpected findings included a cluster in *Macrophomina phaseolina,* that matched the PKS BGC for chaetoglobosins, a class of mycotoxins with both antifungal and anti-cancer activities (Ali, Caggia et al. 2015; Jiang, Song et al. 2017) found in the distantly related *Chaetomium globosum* (Sordariomycetes) and some Eurotiomycetes (Schumann and Hertweck 2007) (Figure 2, Table S1). Another unexpected finding was the similarity between a CO-OCCUR cluster in *M. phaseolina* and the BGC for leucinostatin, a peptaibol compound with putative antimicrobial and antifungal activity, that was previously only known from taxa in the Sordariomycetes (Wang, Liu et al. 2016).

### Cluster co-occurrence networks reveal contrasting trends in diversification

We visualized all significant homolog group co-occurrences predicted by CO-OCCUR as networks where nodes represent homolog groups and edges connect homolog groups that co-occur with unexpected frequency in genomic regions containing core biosynthetic genes (Figure 3a). A total of 33 discrete networks were recovered, with 71% of homolog groups located in the largest two networks. Signature genes tended to be highly connected to other homolog groups in two qualitatively different types of subnetworks. In one type of subnetwork, signature genes are centrally connected to diverse accessory homolog groups (e.g. PKS subnetworks), while in the other type one or more signature genes are non-centrally linked with fewer accessory homolog groups (e.g., the NRPS and DMAT subnetwork in network 1). By quantifying the betweenness centrality of each node (a function of the number of shortest network paths that pass through that node) within each network, we identified signature genes and several other biosynthetic enzymes, transporters, and DNA binding proteins that bridge alternate subnetworks (Figure 3a,b).

**Figure 3.**
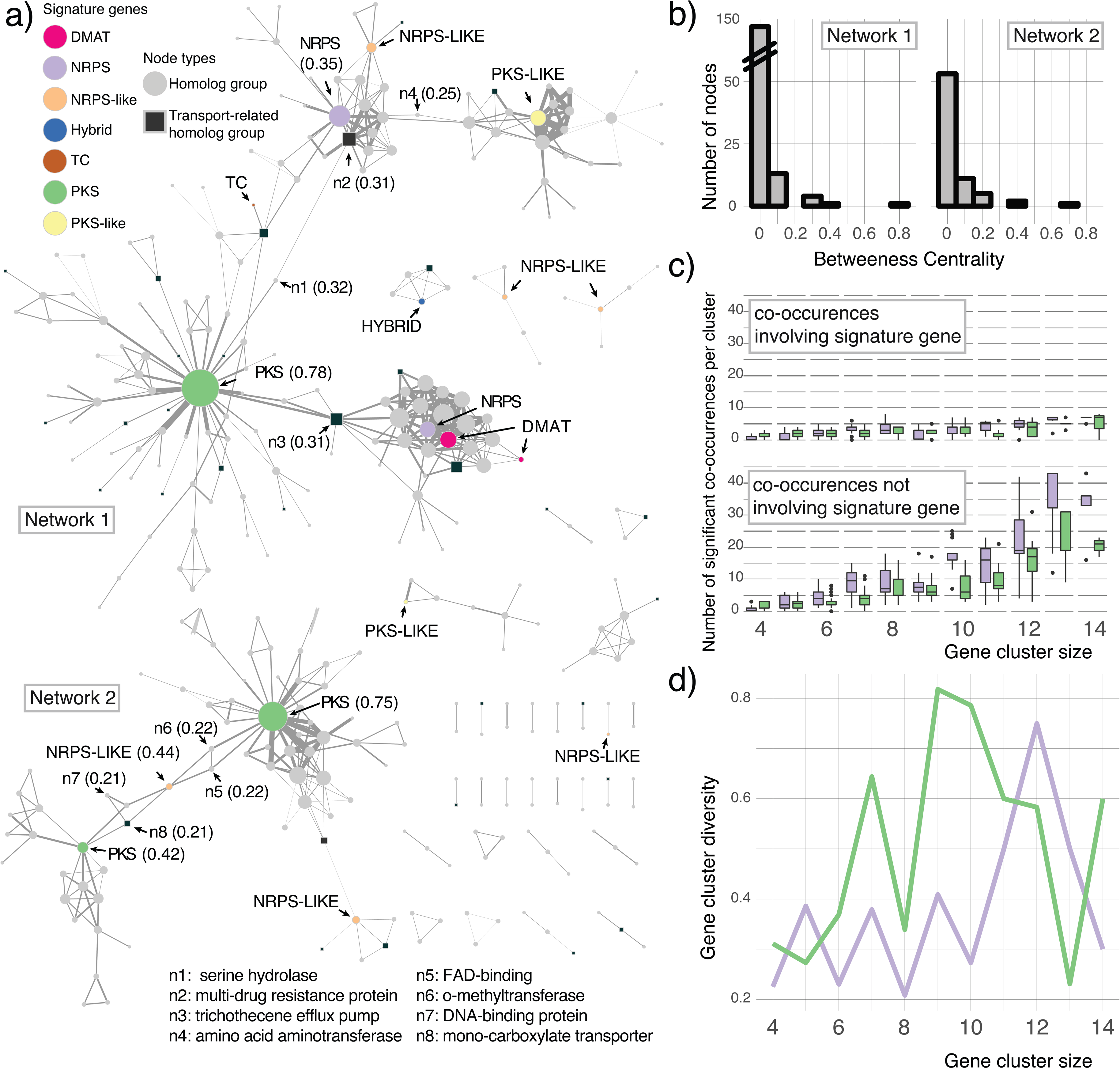
Gene co-occurrence networks among biosynthetic signature gene clusters. **a)** *Co-occurrence network of gene homolog groups (homolog groups).* Nodes in the co-occurrence network represent all homolog groups found in homologous cluster groups (cluster groups). Edges represent significant co-occurrences between homolog groups. Node size is proportional to the number of significant co-occurrences involving that homolog group, and edge width is proportional to the number of unique cluster types (either cluster groups or orphan clusters) with ≥ 4 homolog groups that contain the co-occurrence. Distance between nodes is proportional to the number of co-occurrences they have in common, adjusted by edge width. Signature genes (colored circles) and transport-related function (squares) are indicated. Betweenness centrality scores ≥0.2 are indicated in brackets for signature genes and eight other nodes (n1-8). Networks 1 and 2 are the two largest networks. b) *Histogram of betweenness centrality scores* for all nodes in Networks 1 and 2 (bin width = 0.1). c) *Significant co-occurrences within PKS and NRPS clusters.* Boxplots of homolog group co-occurrences involving signature genes (top) and non- signature genes (bottom) across all polyketide synthase (PKS; green) and nonribosomal polypeptide synthetase (NRPS; purple) clusters with ≥4 unique homolog groups. Boxplots display the 75% percentile (top hinge), median (middle hinge), the 25^th^ percentile (lower hinge), and outliers (dots) determined by Tukey’s method. d) *Diversity of PKS and NRPS clusters*. A line chart tracks the diversity of PKS and NRPS clusters across all cluster sizes for both PKS (green) and NRPS (purple) clusters, where diversity is defined as the total number of unique cluster types (either cluster group or orphan clusters) divided by the total number of clusters.

PKS BGCs are more compositionally diverse than NRPS BGCs. BGCs containing PKS signature genes tended to have fewer significant co-occurrences among their constituent genes across various BGC sizes, compared to BGCs containing NRPS signature genes (Figure 3c). This is consistent with a trend in which clusters containing PKS signature genes have more unique types of BGCs for a given cluster size (corrected by the total number of BGCs of that size), compared with BGCs containing NRPS signature genes (Figure 3d).

### Different algorithms annotate overlapping and complementary sets of clustered genes

CO-OCCUR predictions and the pHMM-based SMURF (Khaldi, Seifuddin et al. 2010) and antiSMASH (Blin, Wolf et al. 2017) programs all predicted similar numbers, but different types of BGCs. antiSMASH identified a total of 1710 clusters that were part of 252 cluster groups and 887 orphan clusters with 4 or more homolog groups (Table SH, Table SI). SMURF identified a total of 686 clusters that were part of 194 cluster groups and 495 orphan clusters with 4 or more homolog groups (Table SJ, Table SK). CO-OCCUR predicted 1469 clusters that are part of 239 cluster groups and 220 orphan clusters with 4 or more homolog groups (Table SC, Table SE). We found that no single algorithm was able to annotate all predicted genes of interest in a BGC, even those predicted to be involved in SM biosynthesis (Figure 4a, Table SL). CO-OCCUR identified 51.2% and 37.7% of the clustered genes detected by SMURF and antiSMASH, respectively. Conversely, SMURF and antiSMASH identify 40.7% and 42.0% of the clustered genes detected by CO-OCCUR, respectively. When examining only genes predicted to participate in SM biosynthesis, transport and catabolism, we found that CO-OCCUR identified 51.2% and 43.3% of genes detected by SMURF and antiSMASH, respectively, while SMURF and antiSMASH each identified 62.6% of those detected by CO-OCCUR (Figure 4a).

**Figure 4.**
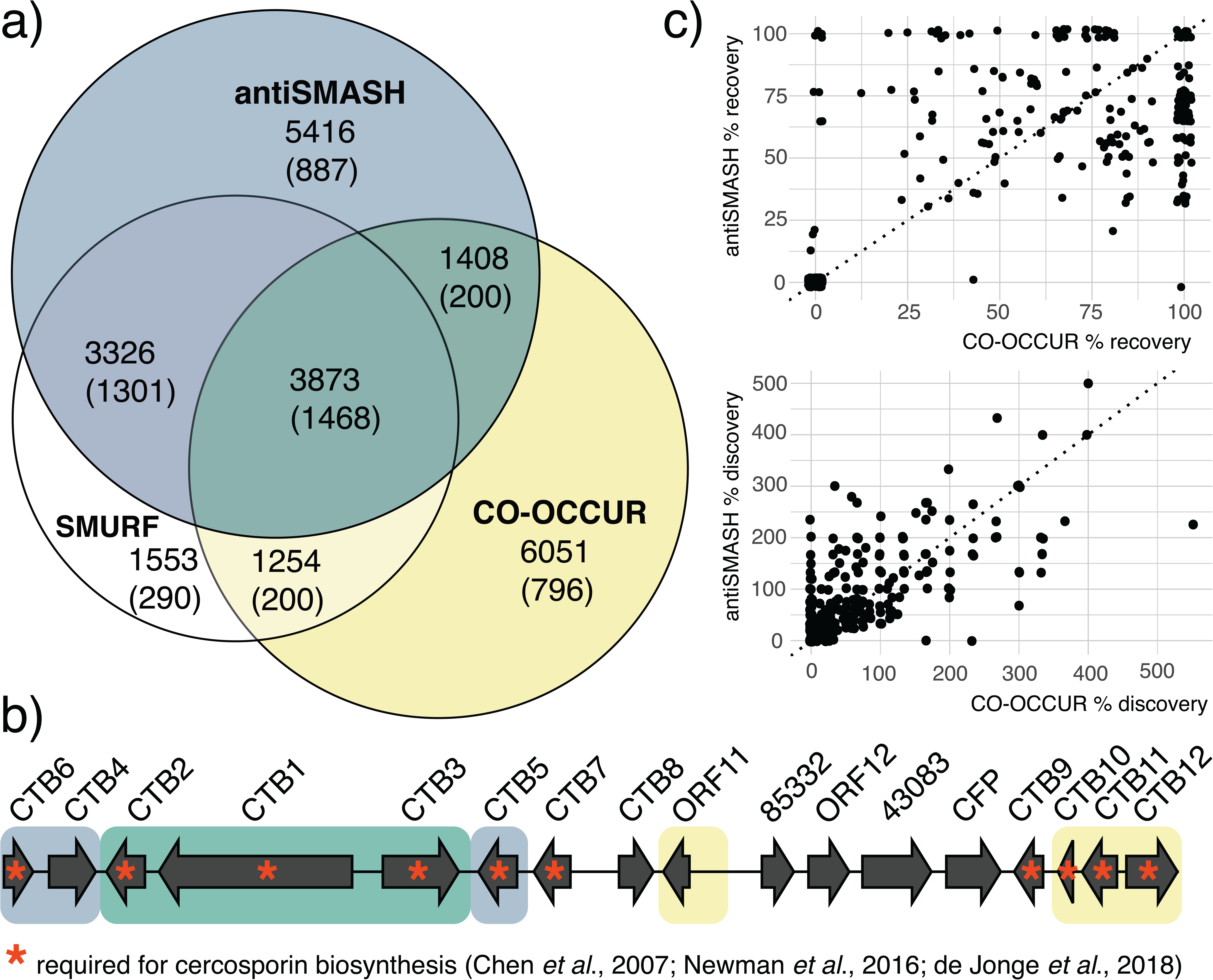
Benchmarking three different algorithms for biosynthetic gene cluster (BGC) detection. *a) Proportional Venn diagram of distinct and overlapping BGC genes of interest detected by SMURF, antiSMASH and CO-OCCUR.* SMURF and antiSMASH use profile Hidden Markov Models (pHMMs) to identify clustered genes of interest, while CO-OCCUR uses linkage-based criteria (see methods). Clustered genes (unbracketed) and secondary metabolism biosynthesis, transport and catabolism clustered genes (fuNOG) detected are indicated for each algorithm/combination. *b) Complementary recovery of the cercosporin BGC using antiSMASH and CO-OCCUR*). Shading of genes in the *Cercospora zeae-maydis* cercosporin BGC (MIBiG ID BGC0001541; recovered clusterID Cerzm1_BGC0001541_h92 in Table SG) indicates genes identified by antiSMASH (blue), CO-OCCUR (yellow), or both algorithms (green). Gene names are as in (de Jonge et al., 2018) and those required for cercosporin biosynthesis (Chen et al., 2007; Newman et al., 2016; de Jonge et al., 2018) are indicated with an asterisk. *c) Gene recovery and discovery in clusters homologous to known BGCs.* Scatterplots show the percent of genes recovered (top) or discovered (bottom) by antiSMASH vs. CO-OCCUR at each locus homologous to a MIBiG BGC (search criteria: minimum 3 gene cutoff; minimum of 75% genes similar to MIBiG BGC genes in locus). Percent recovery is defined as the number of genes identified by BLASTp in an algorithm-identified cluster divided by the size of the BLASTp identified BGC, multiplied by 100. Percent discovery is defined as the number of genes identified by the cluster algorithm but not identified in the BLASTp search, divided by the size of the BLASTp identified BGC, multiplied by 100. y = x at the dotted reference line.

The complementary nature of the CO-OCCUR and antiSMASH algorithms is illustrated by their annotations of a characterized BGC that encodes the biosynthesis of cercosporin, a non-host specific polyketide produced by *Cercospora* spp. (Dothideomycetes) and *Colletotrichum* (Sordariomycetes) (de Jonge, Ebert et al. 2018). Encoded in a BGC, all 10 genes involved in cercosporin biosynthesis, are known and characterized (CTB1-3, CTB5-7, CTB9), in addition to a regulator (CTB8) and two transporters (CTB4 and CFP)(Chen, Lee et al. 2007; de Jonge, Ebert et al. 2018; Newman & Townsend 2016). At this BGC’s locus in *Cercospora zeae-maydis*, both antiSMASH and CO-OCCUR annotated CTB1, CTB2, and CTB3 as genes of interest; only antiSMASH annotated CTB4, CTB5 and CTB6; only CO-OCCUR annotated CTB10, CTB11 and CTB12; and no algorithm annotated CTB7, CTB8, CTB9 or CFP (Figure 4b).

CO-OCCUR and antiSMASH recovered similar proportions of loci homologous to known BGCs and predicted additional genes of interest in the vicinity of these candidate. Using BLASTP, we identified 364 BGCs with ≥ 3 genes across all Dothideomycete genomes that are homologous to 58 characterized BGCs from the MIBiG database (i.e., where ≥75% of genes show similarity) (Table SM). We then compared how many genes within and around these BGCs were predicted to be of interest by either antiSMASH or CO-OCCUR by cross-referencing all BGCs detected by each method, and found that both algorithms recovered similar percentages of BGC content (antiSMASH mean percent recovery = 48.3%, SD = 37.6%; CO-OCCUR mean percent recovery = 51.0%, SD = 42.6%), although for any given BGC, percent recovery often differed between each algorithm (Figure 4c, Table SG). We also found that both antiSMASH and CO-OCCUR identified similar numbers of new genes of interest around BGC loci (antiSMASH mean percent discovery = 65.4%, SD = 85.4%; CO-OCCUR mean percent discovery = 56.6%, SD = 89.4%), and that the number of additional genes of interest often exceeded the size of the recovered candidate cluster. High rates of novel gene discovery are perhaps expected given that many of the clusters in MIBiG are only partially annotated.

### Some over-dispersed clusters have ecologically biased distributions

We found that nearly one-fifth (18%) of cluster groups are phylogenetically over-dispersed when compared to expected distributions that would result from strict Brownian evolution using Fritz and Purvis’ D statistic, where more closely related species are predicted to be more similar to each other compared to more distantly related species (Figure 5, Figure SD). Six over-dispersed cluster groups were over-represented (present at least twice as often) in either plant pathotrophs or plant saprotrophs (Figure 5). By comparison 22.5% of cluster groups had distributions that were more conserved than expected. The remaining cluster group distributions either fell on a continuum between phylogenetically conserved and over-dispersed (35.1%) or were present in sets of taxa too small to be analyzed (23.8%). Figure 6 presents three examples of closely related cluster groups that vary in their phylogenetic conservation (See Methods). Cluster groups in the first group, which partially encode the 1,8-dihydroxynaphthalene (DHN) melanin pathway, were found in nearly all Dothideomycetes; cluster groups in the second group were restricted to the Pleosporales; and cluster groups in the third group were found among *Bipolaris* and *Dothidotthia,* two closely related genera within the Pleosporales.

**Figure 5.**
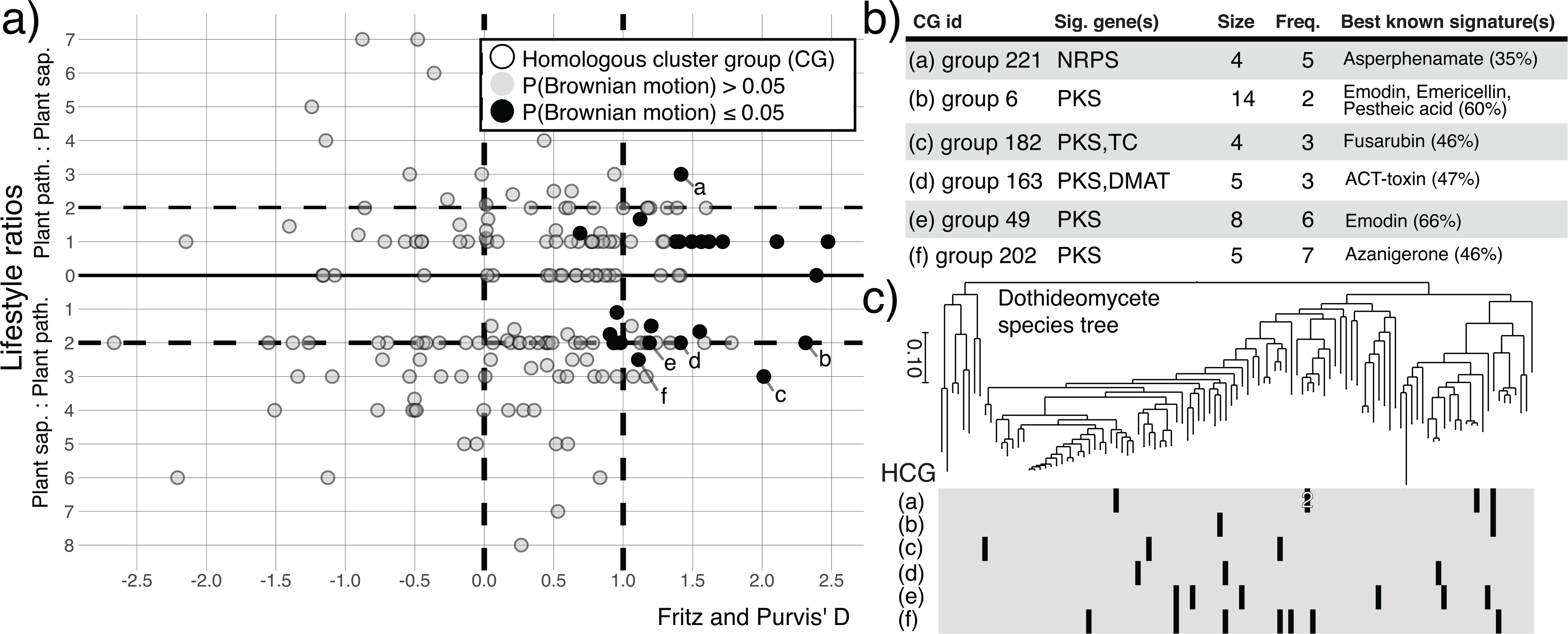
Phylogenetic and ecological signal in the distributions of homologous cluster groups (cluster groups). a) *Scatterplot of phylogenetic and ecological signal of cluster groups*. Values along the X-axis correspond to Fritz and Purvis’ D statistic, representing phylogenetic signal in a cluster group’s distribution. Distributions of cluster groups with D<0 are more conserved compared to a Brownian model of trait evolution, and distributions of cluster groups with D>1 are considered over-dispersed. Cluster groups with more pathotrophs than saprotrophs have Y >0 while cluster groups with more saprotrophs than pathotrophs have Y <0. Point representing cluster group distributions with probability ≤0.05 of Brownian trait evolution are in black, while those >0.05 are in gray. Cluster groups with P(Brownian) ≤0.05 and a lifestyle ratio ≥2 are labeled and described in b) and c). Only cluster groups with ≥ 4 unique gene homolog groups (homolog groups) per cluster are shown. b) *Summary descriptions of labeled cluster groups.* Sig. genes = signature genes present in the cluster group; Size = number of unique homolog groups in the cluster group reference cluster; Freq. = number of fungi with a cluster that belongs to the cluster group; Best known signature(s) = signature gene(s) from the MIBiG database with the highest similarity to signature genes from the cluster group, with average percentage similarity shown in parentheses. c) *Phylogenetic distributions of labeled cluster groups*. Presence (black cells) and absence (gray cells) matrix of clusters assigned to each labeled cluster group across Dothideomycetes genomes tree as in Figure 2.

**Figure 6.**
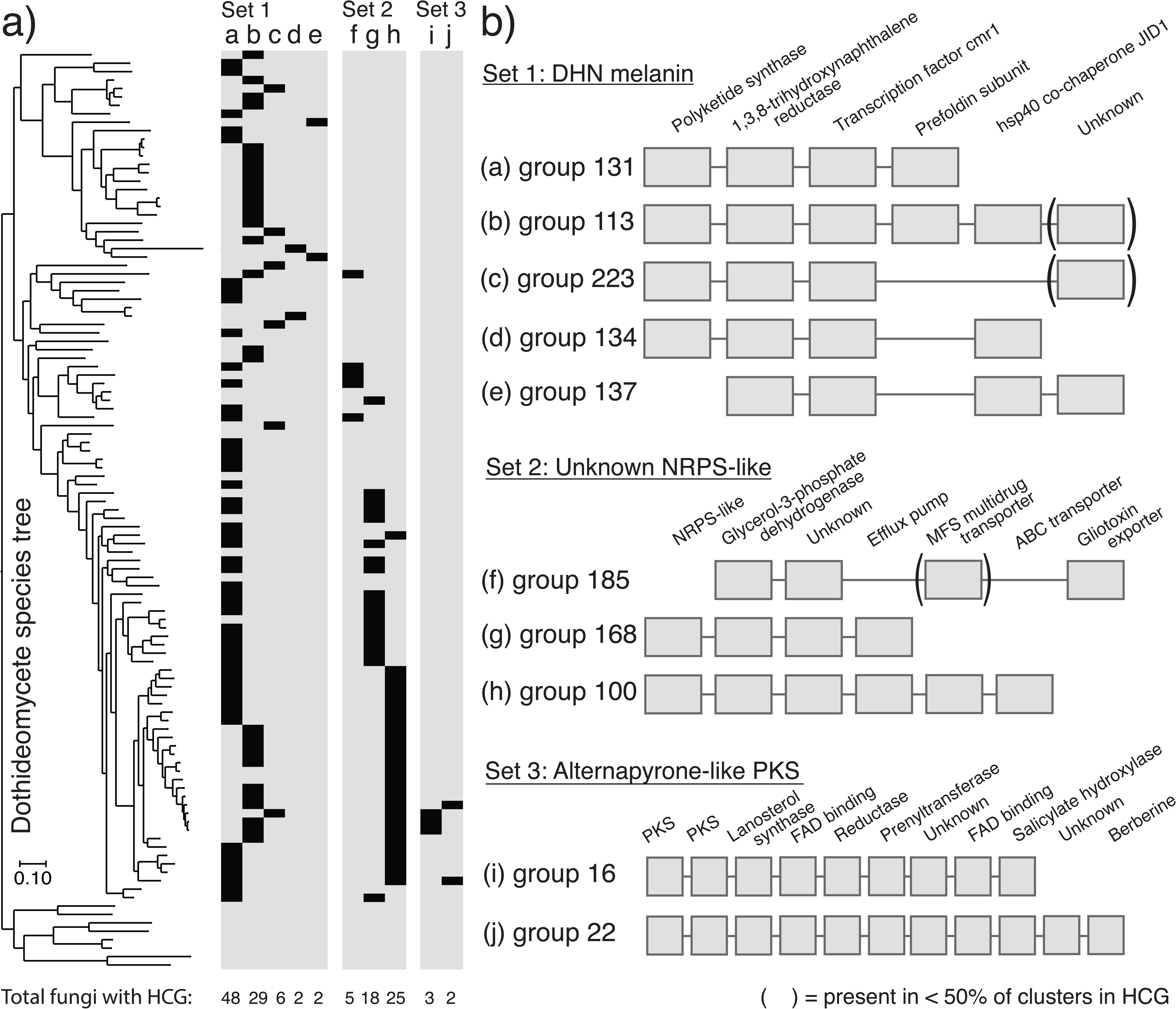
Three examples of homologous cluster groups (cluster groups) with conserved phylogenetic distributions. a) *Cluster group distributions.* Presence (black cells) and absence (gray cells) matrix of clusters assigned to various cluster groups (columns a-j, described in part b), across Dothideomycetes genomes tree as in Figure 2. Each matrix contains distinct sets of cluster groups that are separated by ≤0.05 distance units on the complete linkage tree in Figure 2. The number of fungi with each cluster group is indicated at the bottom of each column. b) *Cluster group composition.* Cluster groups in set 1 are predicted to encode DHN melanin biosynthesis; set 2 contains unknown cluster groups with NRPS-like signature genes; set 3 contains unknown cluster groups with PKS signature genes, where the PKSs from group 16 are on average 84% similar to the PKS in the characterized alternapyrone cluster (MIBiG ID: BGC0000012). Homolog group presence in a given cluster group is indicated by a gray box below the description. Brackets surround homolog groups present in <50% of clusters assigned to a given cluster group.

### Dothideomycetes have five distinct types of DHN melanin clusters

We detected five cluster groups with distinct but overlapping compositions that appear to encode partial pathways for 1,8-dihydroxynaphthalene (DHN) melanin biosynthesis in 87 of the 101 genomes (Figure 6). No genome had more than one predicted DHN melanin cluster. The two most prevalent types, cluster group 131 and cluster group 113, are found in 48 fungi from 10 of the 13 taxonomic orders and in 29 fungi from 6 orders, respectively. Cluster groups 131, 113, 223 and 134 encode 2 of the 5 biosynthetic genes (pks1, 1,3,8-trihydroxynapthalnene [T3HN] reductase) and the transcription factor (cmr1) involved in DHN melanin biosynthesis, while cluster group 137 encodes 1 biosynthetic gene (T3HN reductase) and cmr1 (Figure 6b). In addition to the homolog groups known to participate in DHN melanin biosynthesis, we detected 3 additional homolog groups (Prefoldin subunit, Heat-shock protein 40 co-chaperone JID1, and a protein of unknown function) that are broadly conserved within DHN melanin clusters but that have no known role in melanin biosynthesis. As an example of how CO-OCCUR is not constrained by *a priori* assumptions of pHMMs, these additional homolog groups were not detected by either antiSMASH or SMURF despite their prevalent linkage to the known biosynthetic genes.

### SM cluster diversity is under-sampled and increases proportionally with total number of genomes in Pleosporales

Cluster repertoires (combinations of cluster groups found within a given genome) differ markedly between fungi from different genera (mean pairwise Sørensen dissimilarity = 0.79, SD = 0.12) and to a lesser extent within a genus (mean = 0.37, SD = 0.13), with dissimilarity increasing linearly with phylogenetic distance across all pairwise species combinations among 49 Pleosporales (y = 0.84x + 0.51; r2 = 0.50), the most well-sampled Dothideomycetes order (Figure 7a, Figure SF). However, given the same level of within-repertoire diversity (i.e., alpha diversity) and total diversity across all repertoires (i.e., gamma diversity), dissimilarity between repertoires (i.e., beta diversity) can result from either nestedness (where some repertoires are subsets of others) or turnover (where no repertoire is a subset of the other), or a combination of the two. When we partitioned total Sørensen dissimilarity between all cluster repertoires (β_SOR_ = 0.969) into its nestedness and turnover components, we found that nearly all of the differences between the cluster repertoires of different genomes were due to turnover (β_SIM_ = 0.96) and not repertoire nestedness (β_SNE_ = 0.008), such that any given cluster repertoire contains a unique combination of clusters (Figure SH). Furthermore, the compositional diversity of gene clusters within a given repertoire (measured as the total branch length on a Raup-Crick linkage tree, Figure 7a) scales linearly with repertoire size (y = 0.49x + 3.92; adj. R^2^ = 0.86), indicating that clusters added to a given repertoire are generally dissimilar to the clusters already present in that repertoire (Figure 7b). Finally, rarefaction analysis of the total number of unique cluster groups and orphan clusters (i.e., cluster richness) detected over increasing numbers of sampled genomes suggests genomes within Pleosporales are under-sampled with respect to BGC diversity, and project substantially more unique cluster types arising from future genome sampling within this order (Figure 7c).

**Figure 7.**
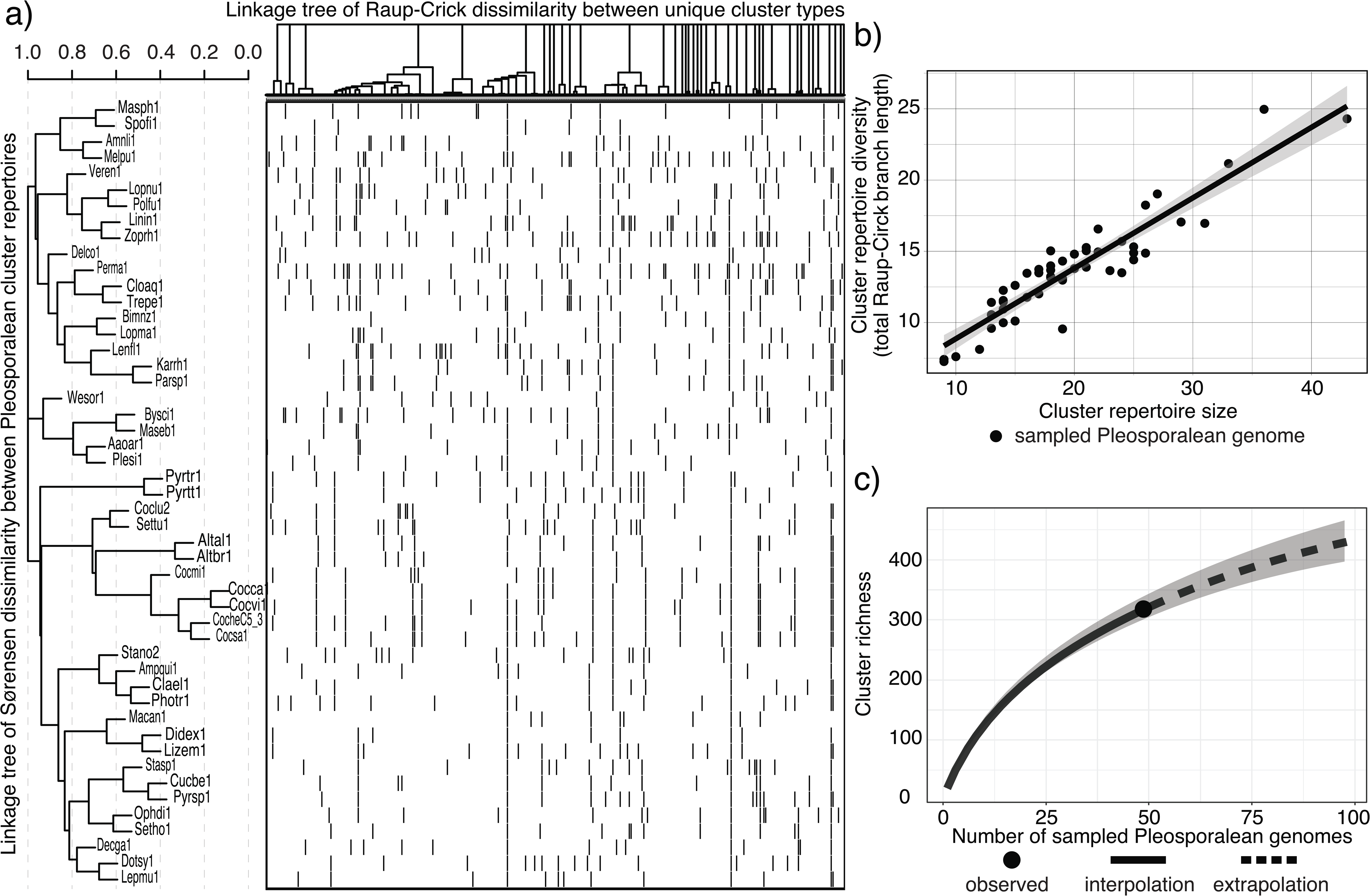
Diversity of secondary metabolite gene cluster repertoires in Pleosporalean fungi. a) *Grouping of fungi based on the combinations of gene clusters (i.e., cluster repertoires) found in their genomes.* Shown to the left is a complete linkage tree where distance between different fungal species is proportional to the Sorensen dissimilarity between their cluster repertoires. To the right is a presence (black) and absence (white) matrix where each column represents a unique cluster type (either a homologous cluster group or cluster orphan) and each row corresponds to the adjacent fungal genome. On top of the heatmap is a complete linkage tree displaying relationships between unique cluster types, where distance is proportional to the Raup-Crick dissimilarity in cluster composition. b) *Relationship between cluster repertoire size and cluster repertoire diversity.* Cluster repertoire diversity was calculated for each genome by finding the total branch length on the Raup-Crick dissimilarity tree in a) associated with the set of clusters found in that genome. Cluster repertoire diversity is thus a measurement of a given genome’s repertoire diversity, in terms of the gene content of its clusters. A solid line models the linear relationship between repertoire size and diversity (adj. R^2^ = 0.855). The shaded area around the line represents the 95% confidence interval associated with the model. c) *Sampled and projected secondary metabolite cluster richness within the Pleosporales.* Rarefied (solid lines) and extrapolated (dotted lines) estimates of secondary metabolite gene cluster richness (i.e., the number of unique cluster types) with respect to the number of sampled genomes are shown for the Pleosporales. Shaded areas represent the 95% confidence intervals for both estimate types, derived from 100 bootstrap replicates. All three graphs were generated using data from the 318 unique cluster types with ≥ 4 unique gene homolog groups that are associated with 47 Pleosporalean fungi and 2 as yet unclassified fungi found within the Pleosporalean clade on the phylogenomic species tree in Figure 2.

## Discussion

BGC diversity has been investigated primarily in bacteria (Cimermancic, Medema et al. 2014) and within individual genera in the fungal classes Eurotiomycetes and Sordariomycetes (Lind, Wisecaver et al. 2017; Theobald, Vesth et al. 2017; Villani, Proctor et al. 2019). Although Dothideomycetes are producers of a number of secondary metabolites important to fungal-plant interactions and toxin production, to date there has not been a systematic evaluation of BGC diversity in the Dothideomycetes nor in any other fungal class. Fungal genomes experience frequent reorganization and changes in gene composition that underlies large-scale differences in chromosomal macro- and micro-synteny among species (Grandaubert, Lowe et al. 2014; Hane, Rouxel et al. 2011; Shi-Kunne, Faino et al. 2018). Yet despite the overall dynamic nature of fungal chromatin, tight linkage is often maintained between loci with related metabolic functions, manifesting as gene clusters (Del Carratore, Zych et al. 2014). Here, we developed an alternative, function-agnostic approach to annotating SM genes of interest that exploits these patterns of microsynteny in order to identify previously unexplored dimensions of fungal BGC diversity.

### Complementary methodologies enhance understanding of BGC composition and diversity in Dothideomycetes

There are two main approaches to predicting genes that are functionally associated in BGCs. The first uses targeted methods based on precomputed pHMMs derived from a set of genes known to participate in SM metabolism to identify sequences of interest (Khaldi, Seifuddin et al., 2010; Blin, Wolf et al., 2017). The second uses untargeted methods based on some function-agnostic criteria, such as synteny conservation or shared evolutionary history, to implicate genes as part of a gene cluster (Gluck-Thaler & Slot 2018). Due to common metabolic functions employed across distantly related taxa, targeted approaches, such as those employed by SMURF and antiSMASH, have proven enormously successful. However, our objective in this study was to develop a complementary untargeted approach in order to capture undescribed BGC diversity within a single fungal lineage.

Our CO-OCCUR algorithm leverages a database of 101 Dothideomycete genomes in order to annotate genes of interest using unexpectedly conserved genetic linkage as an indicator of selection for co-inheritance with SM signature genes. CO-OCCUR failed to recover many of the genes annotated using the pHMM approaches employed by SMURF and antiSMASH, indicating that it has limitations in its prediction of secondary metabolite BGC content. These results suggest that it is not optimal for the *de novo* BGC annotation of individual genomes, and its ability to annotate genes of interest is proportional to their co-occurrence frequency in a given database, meaning that it is not well suited for recovering associated SM genes that are not evolutionarily conserved. This may explain in part why 10,295 genes (including 2,478 genes predicted to be involved in secondary metabolism) identified by antiSMASH and SMURF combined were not detected with CO-OCCUR (Figure 4), and why CO-OCCUR detected only a few of the host-selective toxins found in Dothideomycetes.

Nevertheless, our method avoids some of the limitations intrinsic to algorithms that employ pHMMs to delineate cluster content. While pHMM-based approaches gain predictive power by leveraging similarities in SM biosynthesis across disparate organisms, they may fail to identify gene families involved in secondary metabolism that are unique to a particular lineage of organisms. For example, SMURF detects accessory SM genes using pHMMs derived from mostly *Aspergillus* (Eurotiomycetes) BGCs (Khaldi, Seifuddin et al., 2010), while antiSMASH v4 and v5 use 301 pHMMs of smCOGs (secondary metabolism gene families) derived from aligning SM-related proteins, of which few are currently from fungi, in order to identify genes of interest in the regions surrounding signature biosynthetic genes (Blin, Wolf et al., 2017). Taxonomic bias introduced by sampling a limited number of BGCs may account for the 6,051 proteins found in BCGs that were identified by CO-OCCUR but not any other algorithm, of which 796 are predicted to participate in secondary metabolism and 617 could not be assigned to a COG category but nevertheless have domains commonly observed in secondary metabolite biosynthetic proteins (e.g., methyltransferase, hydrolase).

A linkage-based approach can also identify non-canonical accessory genes involved in SM biosynthesis. For example, we detected 3 genes among the 5 variants of the DHN melanin cluster that were not previously considered to be part of this BGC and not detected by either antiSMASH or SMURF. One of these genes, a predicted HSP40 chaperone, is a homolog of the yeast gene JID1, whose knock-out mutants display a range of phenotypes (https://www.yeastgenome.org/locus/S000006265/phenotype) related to melanin production, including increased sensitivity to heat and chemical stress. We propose that natural selection (not genetic hitchhiking) is responsible for conservation of synteny in these loci, because SM cluster locus composition and microsynteny in general are typically highly dynamic in fungi (Lind, Wisecaver et al., 2018; Proctor, McCormick et al. 2018), and therefore conserved linkage in these clusters over speciation events is a strong indicator of related function (de Jonge, Ebert et al. 2018; Del Carratore, Zych et al. 2019). The identification of genes with non-canonical functions, including those not participating directly in SM biosynthesis, may reveal SM supportive functions, including mechanisms to protect endogenous targets of the metabolic product (Keller 2015), in addition to novel biosynthetic genes (de Jonge, Ebert et al. 2018).

Ultimately, targeted and untargeted approaches to BGC annotation reinforce and enrich our understanding of BGC diversity, as no single method identifies all accessory genes of interest in the regions surrounding signature biosynthetic genes (Figure 4). It is notable that the cercosporin BGC was long thought to consist only of CTB1-8, based on functional analyses and structural prediction. However, de Jonge *et al*. recently predicted CTB9-12 to be of interest after observing that these genes have conserved synteny among all fungi that possessed CTB1-8, and subsequently demonstrated they are essential for cercosporin biosynthesis (de Jonge, Ebert et al. 2018). Only CO-OCCUR detected these additional four genes and bBoth pHMM-based models and CO-OCCUR were required to detect the complete cercosporin BGC in our study. Given the complementary nature of the advantages and disadvantages of different algorithms, we suggest future studies incorporate multiple lines of evidence from both targeted and untargeted approaches to more fully capture BGC compositional diversity. The 332 homolog groups of interest that we identified using CO-OCCUR could further be used to build pHMMs and be incorporated into existing BGC annotation pipelines in order to facilitate more complete analyses of single genomes.

### Signature genes differ in mode of BGC diversification

Although BGCs in fungi typically display characteristics of diversification ‘hotspots’, showing elevated rates of gene duplication and gene gain and loss (Wisecaver, Slot et al. 2014; Lind, Wisecaver et al. 2017), modular parts of clusters and even entire clusters are often shared between divergent species. BGC diversification through gain and loss of individual genes and sub-clusters of genes has been demonstrated in bacterial BGC diversity (Cimermancic, Medema et al. 2014). Although the extent of sub-clustering in fungal genomes has never been directly addressed to our knowledge, the algorithm we designed here essentially functions by identifying the smallest possible type of sub-cluster: a pair of genes found more often than expected by chance. The unexpected co-occurrence of gene pairs revealed that the two largest types of signature gene families, PKS and NRPS, have contrasting co-occurrence network structures. NRPS homolog groups are embedded in highly reticulate cliques (i.e., form unexpected associations with genes that co-occur amongst themselves). This could suggest NRPS cluster diversification is constrained by interdependencies among accessory genes. By contrast, PKS homolog groups are network hubs (i.e., form unexpected associations with many non-co-occurring genes), which may underlie the higher compositional diversity and decreased frequency of unexpected co-occurrences found within PKS clusters (Figure 3c, d). The apparent contrast in how these different signature cluster types are assembled may reflect the range of accessory modifications typically applied to the structures of polyketides and nonribosomal peptides produced by PKSs and NRPSs. Alternatively, PKS clusters may be subject to more diversifying selection, due to the ability of cognate metabolism in other organisms to utilize, degrade, or neutralize the metabolites. These hypotheses remain to be tested.

### Persistent gene co-occurrences reveal layers of combinatorial evolution

Previous large-scale analyses of BGCs suggest there is an upper limit to the number of gene families that associate with signature biosynthetic genes, and that diversity is in large part dependent on combinatorial re-shuffling of existing loci (Cimermancic, Medema et al. 2014). Our analysis expands the number of gene families implicated in BGC diversity and identifies patterns of modular combinatorial evolution among accessory homolog groups with metabolic, transport and regulatory-related functions. While some of these accessory homolog groups are restricted to BGCs with a particular type of signature SM gene, others are present in multiple BGC types, suggesting they encode evolvable or promiscuous functions that can be readily incorporated into different metabolic processes (Table SB). For example, 34 homolog groups with predicted transporter functions are common features of the clusters we detected, present in just under half (43%) of all predicted clusters. Among these homolog groups, 5 have been recruited to compositionally diverse gene clusters and are primarily annotated as toxin efflux transporters or multidrug resistance proteins. Transporters are a key component of fungal chemical defense systems, well known for facilitating resistance to fungicides and host-produced toxins (Coleman & Mylonakis 2009). Transporters are also increasingly recognized as integral components of self-defense mechanisms against toxicity of endogenously produced SMs (Menke, Dong et al., 2012).

### Heterogeneous dispersal patterns of BGCs underpin fungal ecological diversity

The distribution of fungal chemodiversity remains difficult to observe and interpret directly, making BGCs useful tools for elucidating underlying trends in fungal chemical ecology. Although the vast majority of BGCs remain uncharacterized, their phylogenetic distributions occasionally provide clues to the selective environments that promote their retention (Slot 2017). For example, spotty distributions resulting from horizontal transfer of BGCs between distantly related but ecologically similar species suggests the encoded metabolites contribute to fitness in the shared environment (Dhillon, Feau et al. 2015; Reynolds, Vijayakumar et al. 2018). Shared ecological lifestyle may also help explain why certain clusters, such as those involved in putative degradative pathways, are retained among phylogenetically distant species (Gluck-Thaler and Slot 2018). Our simple eco-evolutionary screen identified 43 BGCs that are more widely dispersed than expected under neutral evolutionary models, and further revealed that a subset of these BGCs are present more often in fungi with specific nutritional strategies (e.g. plant saprotrophs and plant pathotrophs), suggesting the molecules they encode contribute to specific plant-associated lifestyles (Figure 5). For example, we found an over-dispersed NRPS BGC (group 221) that is present in three plant pathogens and one plant saprotroph. In contrast, the 54 BGCs showing a phylogenetically under-dispersed distribution among mostly closely related genomes is consistent with lineage-favored traits, which may or may not be due to shared ecology. For example, a monophyletic clade of 26 pleosporalean fungi all have a 6 gene NRPS-like cluster (group 100) of unknown function, fully maintained among these allied taxa (and a single distant relative), suggesting it encodes a trait that contributes to the success of this lineage. Phylogenetic screens, especially when coupled with more robust phylogenetic analyses, will be useful for prioritizing the characterization of BGCs most likely to contribute to the success of particular guilds or clades.

Among those BGCs with hits to the MIBiG database, we identified clusters that displayed both lineage specific and spotty or sporadic distributions. The Pleosporales, for example, contains many plant pathogens and the conservation of BGCs involved in production of general virulence factors towards plants such as solanopyrone, alternapyrone, and the extracellular siderophore dimethylcoprogen across many taxa in this order suggests a shared lineage-specific trait with roles in plant-pathogenesis. In contrast, the aflatoxin-like cluster Dothistromin cluster, which was proposed to be horizontally transferred from *Aspergillus* (Eurotiomycetes), had a very spotty distribution, found only in several closely related taxa in Capnodiales, supporting a hypothesis of HGT. Similarly, the ETP toxin sirodesmin shares 6 genes with the BGC producing the epipolythiodioxopiperazine (ETP) toxin gliotoxin, which plays a role in virulence towards animals in human pathogen *Aspergillus fumigatus* (Eurotiomycetes) (Gardiner, Cozijnsen et al. 2004; Bok, Chung et al. 2006). Related ETP-like BGCs have since been identified in a number of other taxa of Eurotiomycetes and Sordariomycetes, but among Dothiodeomycetes were previously known only from *L. maculans* and a partial cluster in *Sirodesmin diversum* lacking the core NRPS (Patron, Waller et al. 2007). We detected homologs of this cluster found sporadically distributed in several other taxa within Pleosporales (Figure 2, Table SG). The CO-OCCUR algorithm detected only a few BGC with hits to host-selective toxins (sirodesmin and T-toxin) but failed to detect several well-known host-selective toxins such as HC-toxin, and other host-selective toxins in *Alternaria* alternata. Either these host-selective toxins are not represented in MIBiG or as discussed above, the uniqueness of these clusters and rarity of the linkages between genes in these clusters in the overall dataset may make them difficult to detect through CO-OCCUR.

### Variation among BGC repertoires is due to high BGC turnover, not nestedness

Recent comparative studies have documented high intraspecific diversity of SM pathways within and between different species of plants, bacteria and fungi (Penn et al., 2009; Choudoir, Pepe-Ranney & Buckley 2018; Holeski, Hillstrom et al. 2012; Holeski, Keefover-Ring et al. 2013; Vesth, Nybo et al. 2018). However, identical estimates of diversity can result from two distinct processes: nestedness, where one set of features is entirely subsumed within another, or turnover, where differences are instead due to a lack of overlap among the features of different sets (Baselga 2012). When we partitioned diversity among BGC repertoires in Pleosporales (i.e., β diversity), we found that the vast majority of variation is due to a high degree of genome-specific cluster combinations, and not nestedness (Figure 7, Figure SH). Much of the turnover in BGC repertoire content between genomes appears to occur over relatively short evolutionary timescales (Figure SF), and then diversifies more gradually, suggesting that divergence in repertoires may be closely linked to speciation processes, such as niche differentiation or geographic isolation. Directional selection, especially for multi-genic traits encoded at a single locus (e.g., BGCs), leads to rapid gain/loss dynamics exemplary of many SM phenotypes and genotypes (Choudoir, Pepe-Ranney & Buckley 2018; Lind, Wisecaver et al. 2017). Niche differentiation further reinforces divergence between closely related repertoires, which might lead to rapid accumulation of variation over short evolutionary timescales. Indeed, evidence from within populations suggests that BGCs are occasionally located in genomic regions experiencing selective sweeps in geographically isolated pathogen populations (Hartmann, McDonald et al. 2018). The retention/loss of certain SM clusters is coincident with speciation in bacteria (Kurmayer, Blom et al., 2015) and much of the variation in cluster repertoires in *Metarhizium* insect pathogens is species specific (Xu, Luo et al., 2016). Within Dothideomycetes, the evolution of host-selective toxins even within a single species of pathogen, for example, may allow for niche differentiation, host specialization, and potentially speciation. Rare chemical phenotypes, especially with regards to defense chemistry, may also increase fitness in complex communities (Kursar, Dexter et al. 2009).

### BGCs interact with dimensions of chemical diversity

Biological activity of a SM can increase organismal fitness, but any given molecule is not likely to be biologically active. The screening hypothesis posits that mechanisms to generate and retain biochemical diversity would therefore be selected, despite the energetic costs, because increasing structural diversity increases the probability of “finding” those that are adaptive (Firn and Jones 2003). This phenomenon is analogous to the mammalian immune system’s latent capacity to generate novel antibodies, resulting in a remarkable ability to respond to diverse antagonists (Firn and Jones 2003). However, while the screening hypothesis may equally apply to plants and microorganisms, patterns of diversity we observe here suggest each lineage generates and maintains biochemical diversity in fundamentally distinct ways. Specifically, fungal individuals appear to maximize total chemical beta-diversity while simultaneously minimizing alpha-diversity of similar chemical classes (Nielsen, Grijseels et al. 2017; Vesth, Nybo et al. 2018). In contrast, individual plants are more likely to produce diverse suites of structurally similar molecules (Li, Bladwin & Gaquerel 2015; Song, Qiao et al. 2017; Weinhold, Ullah et al. 2017). We show that total cluster diversity increases linearly with repertoire size across a broad sample of fungi, extending previous observations that individual fungal genomes are streamlined to produce molecules that share little structural similarity. Rather than maintaining sets of homologous BGCs and pathways within the same genome, evidence from ours and other studies suggests that fungi instead maintain high genetic variation in homologous BGCs across individuals at the level of the pan-genome (Ziemert, Lechner et al. 2014; Lind, Wisecaver et al. 2017; Olarte et al. 2019). Although not a selectable evolvability mechanism per se, greater access to the diversity of BGCs harbored in pan-genomes through recombination, hybridization and horizontal transfer effectively outsources the incremental screening for bioactive metabolites across many individuals, thereby decreasing the costs for generating diversity for any given individual and likely accelerating the rate at which effective bioactive metabolite repertoires are assembled within a given lineage (Slot and Gluck-Thaler 2019). Our characterization of BGC diversity across the largest fungal taxonomic class represents a step towards elucidating the broader consequences of these contrasting strategies for generating and maintaining biodiversity of metabolism writ large.

## Conclusions

Fungi produce a range of secondary metabolites that are linked to different ecological functions or defense mechanisms, playing a role in adaptation over time. Although studied at intra- and interspecific level, this phenomenon has not been studied at macroevolutionary scales. The Dothideomycetes represent the largest and phylogenetically most diverse class of fungi, displaying a range of fungal lifestyles and ecologies. Here we assessed the patterns of diversity of biosynthetic gene clusters across the genomes of 101 Dothideomycetes to dissect patterns of evolution of chemodiversity. Our results suggest that different classes of BGCs (e.g. PKS versus NRPS) have differing diversity of cluster content and connectedness among networks of co-occurring genes and implicate high rates of BGC turnover, rather than nestedness, as the main contributor to the high diversity of BGCs observed among fungi. Consequently, little overlap was found in biosynthetic gene clusters from different genera, consistent with diverse ecologies and lifestyles among the Dothideomycetes, and suggesting that most of the metabolic capacity of this fungal class remains to be discovered.

## Methods

### Dothideomycetes genome database and species phylogeny

A database of 101 Dothideomycetes annotated genomes, gene homolog groups, and the corresponding phylogenomic species tree were obtained from (Grigoriev et al. 2014; Haridas et al *in press*).

### Gene cluster annotation with the SMURF algorithm

We used a command-line Python script based on the SMURF algorithm (Vesth, Nybo et al. 2018). Using genomic coordinate data and annotated PFAM domains of predicted genes as input, the algorithm predicts seven types of SM clusters based on the multi-PFAM domain composition of known ’backbone’ genes. The cluster types are 1) Polyketide synthases (PKSs), 2) PKS-like, 3) nonribosomal peptide-synthetases (NRPSs) 4) NRPS-like, 5) hybrid PKS-NRPS, 6) prenyltransferases (DMATS), and 7) terpene cyclases (TCs). The borders of clusters are determined using PFAM domains that are enriched in characterized SM clusters, allowing up to 3 kb of intergenic space between genes, and no more than 6 intervening genes that lack SM-associated domains. SM-associated PFAM domains were borrowed from Khaldi et al. (2010).

### Gene cluster annotation with antiSMASH

All genomes were annotated using antiSMASH v4.2.0 by submitting genome assemblies and GFF files to the public web server with options “use ClusterFinder algorithm for BGC border prediction” and “smCOG analysis” (Blin, Wolf et al., 2017). antiSMASH reports all genes within the borders of a predicted cluster as part of the cluster. For our analysis, we only considered genes belonging to annotated smCOGs or signature biosynthetic gene families as part of a given cluster and excluded all others, in order to obtain conservative, high confidence estimations of cluster content based on genes of interest.

### Sampling null homolog group-pair distributions

We created null distributions from which we could empirically estimate co-occurrence probabilities by randomly sampling homolog group pairs without replacement from all Dothideomycete genomes (Figure SA). Before beginning, we defined null distributions based on two parameters: a range of sizes for the smallest homolog group in the pair, and a range of sizes for the largest homolog group in the pair, where each range progressively incremented by 25 from 1-800 and all combinations of ranges were considered. For example, there existed a null distribution for homolog group pairs where the smallest homolog group had between 26-50 members, and the largest homolog group had between 151-175 members. To begin, we randomly sampled a genome and then randomly selected two genes within 6 genes of one other from that genome. We retrieved the homolog groups to which those genes belonged, and then counted the number of times members of each homolog group were found within 6 genes of each other across all Dothideomycete genomes. We counted the number of members belonging to each homolog group, excluding those that were found within 6 genes of the end of a contig, in order to obtain a corrected size for each homolog group that accounted for variation in assembly quality. The co-occurrence observation was then stored in the appropriate null distribution based on the corrected sizes of each homolog group. For example, the number of co-occurrences of a sampled homolog group pair where the smallest homolog group had a corrected size of 89, and the largest homolog group had a corrected size of 732 would be placed in the null distribution where the smallest size bin was 76-100 members, and the largest size bin was 726-750. All homolog groups with greater than 800 members were assigned to the 776-800 size bin. This sampling procedure was repeated 500,000 times. After evaluating various bin sizes, we ultimately decided to use a range of 25 because this resulted in the most even distribution of samples across all null distributions. Due to variation in the number of homolog groups with any given size across our dataset, it was not possible for all null distributions to contain the same number of samples.

### The CO-OCCUR pipeline

Current BGC detection algorithms first identify signature biosynthetic genes using profile Hidden Markov Models (pHMMs) of genes known to participate in SM biosynthesis, and then search predefined regions surrounding signature genes for co-located “accessory” biosynthetic, regulatory, and transport genes. The approach of CO-OCCUR, in contrast, is to define genes of interest based on whether they are ever found to have unexpectedly conserved syntenic relationships with other genes in the vicinity of signature biosynthetic genes, agnostic of gene function. Here, we used CO-OCCUR in conjunction with a preliminary SMURF analysis to arrive at our final BGC annotations (Figure SA). We first took all SMURF BGC predictions and extended their boundaries to genes within a 6 gene distance that belonged to homolog groups found in another SMURF BGC, effectively “bootstrapping” the BGC annotations in order to ensure consistent identification of BGC content across the various genomes. SMURF BGCs at this point in the analysis were considered to consist of all genes found within the cluster’s boundaries. For each pair of genes in each BGC (including signature biosynthetic genes), we retrieved their homolog groups, and kept track of how many times that homolog group pair was observed across all BGCs. Then, for each observed homolog group pair, we divided the number of randomly sampled homolog group pairs in the appropriate null distribution (based on the corrected sizes of the smallest and largest homolog groups within the observed pair, see above) that had a number of co-occurrences greater than or equal to the observed number of co-occurrences by the total number of samples in the null distribution. In doing so, we empirically estimated the probability of observing a homolog group pair with at least that many co-occurrences by chance, given the sizes of the homolog groups. In this way, we were able to take into account the relative frequencies of each homolog group within a pair across all genomes when assessing the probability of observing that pair’s co-occurrence. For example, if we observed that homolog group 1 and homolog group 2 co-occurred 19 times within SMURF-predicted BGCs, and that homolog group 1 had 57 members while homolog group 2 had 391 members, we would count the number of randomly sampled homolog group pairs that co-occurred 19 or more times within the null distribution where the smallest homolog group size bin was 51-75 and the largest homolog group size bin was 376-400, and then divided this by the total number of samples in that same null distribution to obtain the probability of observing homolog group 1 and homolog group 2’s co-occurrences by chance. All co-occurrences with an empirical probability estimate of ≤0.05 were considered significant and retained for further analysis. In order to decrease the risk of false positive error, we did not evaluate the probability of observing any homolog group pairs with less than 5 co-occurrences, and also did not evaluate any homolog group pairs whose corresponding null distribution had fewer than 10 samples.

Next, in order to obtain our final set of predicted BGCs, we took all homolog groups found in significant co-occurrences, and conducted a *de novo* search in each genome for all clusters containing genes belonging to those homolog groups within a 6 gene distance of each other. In this way, all BGC clusters in our final set consisted of genes that belonged to these homolog groups of interest, while all other intervening genes were not considered to be part of the cluster. We treated homolog groups containing signature biosynthetic genes as we would any other homolog group: if a signature gene predicted by SMURF was not a member of a homolog group part of an unexpected co-occurrence, we did not consider it part of any clusters. We stress that co-occurrences were only used to determine homolog groups of interest, but that once those homolog groups were identified, they did not need to be part of an unexpected co-occurrence within a predicted cluster in order to be considered part of that cluster. By focusing only on genes that form unexpected co-occurrences, it is likely that we have underestimated the compositional diversity of Dothideomycetes BGCs (but this may be the case for all cluster detection algorithms; see Results).

We then grouped all predicted BGCs into homologous cluster groups (cluster groups) based on a minimum of 90% similarity in their gene content, rounded down, in order to obtain a strict definition of BGC homology that increases the likelihood that homologous clusters encode similar metabolic phenotypes. This meant that clusters with sizes ranging from 2-10 were allowed to differ in at most 1 gene; clusters with sizes ranging from 11-20 were allowed to differ in at most 2 genes, etc. Clusters that were not at least 90% similar to any other cluster in the dataset were designated orphan clusters. Note that because there is no perfect way to determine homology when using similarity based metrics, (e.g., a 10 gene cluster could be 90% similar to a 9 gene cluster, which in turn could be 90% similar to a 8 gene cluster, but that 8 gene cluster cannot be 90% similar to the 10 gene cluster), we developed a heuristic approach for sorting clusters into groups. First, we conducted an all-vs-all comparison of content similarity to sort all clusters into preliminary groups by iterating through the clusters from largest to smallest, where size equaled the number of unique homolog groups, and clusters could only be assigned to a single group. Then, within each preliminary group, we identified clusters most similar to all other clusters within the group and used them as references to which all other clusters were compared during a new round of group assignment. In this final round, clusters were grouped together with a given reference into a cluster group if they were at least 90% similar to it and were classified as orphan clusters if they were not 90% similar to any references. The often-unique compositions of clusters means that in most cases, there is no ambiguity to how the clusters are classified; however, for a small number of clusters, especially those with fewer genes, there may be some ambiguity as to which group they belong.

### Annotation of BGCs and gene functions

In order to detect loci homologous to known BGCs in Dothideomycete genomes, amino acid sequences of each annotated BGC within the MIBiG database (v1.4) were downloaded and used as queries in a BLASTp search of all Dothideomycete proteomes (last accessed 04/01/2019). All hits with ≥50 bitscore and ≤1x10^-4^ evalue were retained, and clusters composed of these hits were retrieved using a maximum of 6 intervening genes. In order to retain only credible homologs of the annotated MIBiG queries and to account for error in BLAST searches due to overlapping hits, we retained clusters with at least 3 genes that recovered at least 75% of the genes in the initial query. This set of high confidence MIBiG BGCs was then compared to the set of BGCs predicted by CO-OCCUR and antiSMASH to assess the ability of each algorithm to recover homologs to known clusters. For each algorithm and each BGC recovered using BLASTp to search the MIBiG database, we calculated percent recovery, defined as the number of genes identified by the BLASTp search that were also identified as clustered by the algorithm, divided by the size of the BGC identified by the BLASTp search, multiplied by 100. We also calculated percent discovery, defined as the number of clustered genes identified by the algorithm but not identified in the BLASTp search, divided by the size of the BGC identified by the BLASTp search, multiplied by 100.

In order to annotate BGCs recovered by CO-OCCUR with characterized clusters, we used amino acid sequences of all signature biosynthetic genes in CO-OCCUR clusters as BLASTp queries in a search of the MIBiG database (min. percent similarity=70%; max evalue=1x10^-4^; min. high scoring pairs coverage=50%). Basing our annotations on percent amino acid similarity to characterized signature biosynthetic genes rather than on the number of genes with similarity to BGC genes enabled a more conservative and comprehensive approach, as many BGC entries within the MiBIG database are not complete.

Proteins within predicted BGCs were annotated using eggNOG-mapper (Huerta-Cepas, Forslund et al., 2017) based on fungal-specific fuNOG orthology data (Huerta-Cepas, Szklarczyk, et al., 2015). Consensus annotations for all homolog groups were derived by selecting the most frequent annotation among all members of the group.

### Comparing BGC detection algorithms

In order to assess the relative performances of SMURF, antiSMASH and CO-OCCUR, we compared all BGCs predicted by each method, and kept track of the genes within those BGCs that were identified by either one or multiple methods. We summarized these findings in a venn diagram using the “eulerr” package in R (Larsson 2019). Note that for the purposes of this analysis, BGCs predicted by SMURF and antiSMASH were considered to be composed only of genes that matched a precomputed pHMM, and BGCs predicted by CO-OCCUR were composed only of genes belonging to homolog groups that were part of unexpected co-occurrences, while all other intervening genes within the BGC’s boundaries were not considered to be part of the cluster. In doing so, we effectively ignored intervening genes that were situated between or are immediately adjacent to these clustered genes of interest for the purposes of defining a cluster’s content. While this approach likely does not capture the full diversity of cluster composition, it is expected to decrease false positive error in BGC content prediction and represents a conservative approach to identifying what genes make up a given cluster.

### Construction of a co-occurrence network

We visualized relationships between homolog group pairs with unexpectedly large numbers of co-occurrences in a network using Cytoscape v.3.4.0 (Shannon, Markiel et al. 2003). The network layout was determined using the AllegroLayout plugin with the Allegro Spring-Electric algorithm. In order to identify hub nodes within the network, we calculated betweeness centrality, a measurement of the shortest paths within a network that pass through a given node, for each node using Cytoscape.

### Assessment of cluster group phylogenetic signal

In order to quantify the dispersion of phylogenetic distributions of cluster groups predicted by CO-OCCUR, we created a binary genome x cluster group matrix for all 239 cluster groups with ≥4 genes that indicated the presence or absence of these cluster groups across all 101 genomes. We used this matrix in conjunction with the “phylo.d” function from the “caper” package v1.0.1 in R (Orme, Freckleton et al. 2012) to calculate Fritz and Purvis’ D statistic for each cluster group’s distribution, where D is a measurement of phylogenetic signal for a binary trait obtained by calibrating the observed number of changes in a binary trait’s evolution across a phylogeny by the mean sum of changes expected under two null models of binary trait evolution. The first null model simulates the phylogenetic distribution expected under a model of random trait inheritance, and the second simulates the phylogenetic distribution expected under a threshold model of Brownian evolution that evolves a trait along the phylogeny under a Brownian process where variation in that trait’s distribution accumulates at a rate proportional to branch length (Fritz & Purvis 2010). D≈1 if the trait has phylogenetically random distribution; D≈0 if the trait has a phylogenetic distribution that follows the Brownian model; D>1 if the trait has a phylogenetic distribution that is less conserved, or over-dispersed, compared to the Brownian model; D < 0 if the trait has a phylogenetic distribution that is more conserved, or under-dispersed, compared to the Brownian model.

### Dissimilarity and Diversity Analyses

We created cluster group and orphan cluster x homolog group matrices in order to determine the dissimilarity between predicted cluster groups. In these matrices, for each cluster group or orphan cluster, we indicated the presence or absence of homolog groups in at least one cluster within the cluster group or orphan cluster, effectively summarizing each cluster group and orphan cluster by integrating over the content of all clusters assigned to that group. We next used the matrix in conjunction with the “vegdist” function from the “vegan” package in R (Oksanen, Blanchet et al. 2016) to create a Raup-Crick dissimilarity matrix that was visualized as a dendrogram using complete linkage clustering as implemented in the “hclust” function from the core “stats” package in R. These dendrograms were then used to assess the functional diversity of BGC repertoires (e.g., in the Pleosporales) by measuring the total branch distance connecting all cluster groups and orphan clusters within a given repertoire using the “treedive” function from the “vegan” package in R.

We used the same above procedure to calculate Sørensen dissimilarity between Pleosporalean genomes based on their BGC repertoires, only this time using a genome x cluster group and orphan cluster matrix that depicted the presence or absence of cluster groups and orphan clusters across all 49 Pleosporalean genomes. We also used this matrix to calculate and partition β diversity in Pleosporalean cluster repertoires using the “beta.sample” function (index.family = “sorensen”, sites = 10, samples = 999) from the “betapart” v1.4 package in R (Baselga & Orme 2012) in order to determine how much of the observed diversity among repertoires was due to gain/loss of cluster groups and orphan clusters, and how much was due to nestedness. We also used the genome x cluster group and orphan cluster matrix to conduct a rarefaction of cluster richness across Pleosporalean genomes using the “iNEXT” function (q = 0, datatype=“incidence_raw”, endpoint=98) from the “iNEXT” package in R (Hseih, Ma & Chao 2016).

## Abbreviations

BGC: Biosynthetic Gene Cluster
pHMM: profile Hidden Markov Model
SM: Secondary Metabolite
PKS: Polyketide Synthetase
NRPS: Nonribosomal Peptide Synthetase
TC: Terpene Cyclase
DMAT: dimethylallyl tryptophan synthase

## Declarations

### Ethics approval and consent to participate

No human subjects were involved in the research

### Consent for publication

No human data was used in the research

### Availability of data and materials

All genome data is available at https://mycocosm.jgi.doe.gov/mycocosm/home and described in Haridas et al. *in press*.

All scripts used in the analyses are available at https://github.com/egluckthaler/co-occur Additional data generated in this study is included in the supplemental datafile

### Authors’ contributions

Formulated the study EG-T, KEB, JCS

Designed the methodology EG-T, JCS

Generated resources EG-T, PWC, IG, MB

Collected the data EG-T, SH

Analyzed the data EG-T

Provided leadership and/or mentorship in the study JWS, JCS

Prepared the manuscript EG-T, JCS

Contributed to the writing/editing of the manuscript - KEB, PWC, JWS

## Acknowledgements

Computational work by EG-T was conducted using the resources of the Ohio Supercomputer Center.

## Supporting information

**Table SA.** Gene homolog groups (homolog groups) part of unexpected co-occurrences.

**Table SB.** Unexpected co-occurrences between gene homolog groups (homolog groups) occurring in the vicinity of signature biosynthetic genes, and their frequency across different SM classes.

**Table SC.** Positional information of all recovered CO-OCCUR clusters.

**Table SD.** Genomes used in this study.

**Table SE.** Cluster types (groups and orphans detected by CO-OCCUR.

**Table SF.** BLAST-based annotation of CO-OCCUR clusters with known signature biosynthetic genes from the MIBiG database.

**Table SG.** Cross-referencing clusters retrieved by CO-OCCUR, antiSMASH, and BLAST searches of MIBiG database to determine percent recovery and discovery.

**Table SH.** positional information of all recovered antiSMASH clusters. **Table SI**. Cluster types (groups and orphans detected by antiSMASH. **Table SJ**. positional information of all recovered SMURF clusters.

**Table SK.** Cluster types (groups and orphans detected by SMURF).

**Table SL.** Overlapping and complementary recovery of clustered genes of interest using antiSMASH, SMURF, and CO-OCCUR.

**Table SM.** Positional information of all clusters recovered with a BLASTp search of the MIBiG database.

---

**Figure SA.**
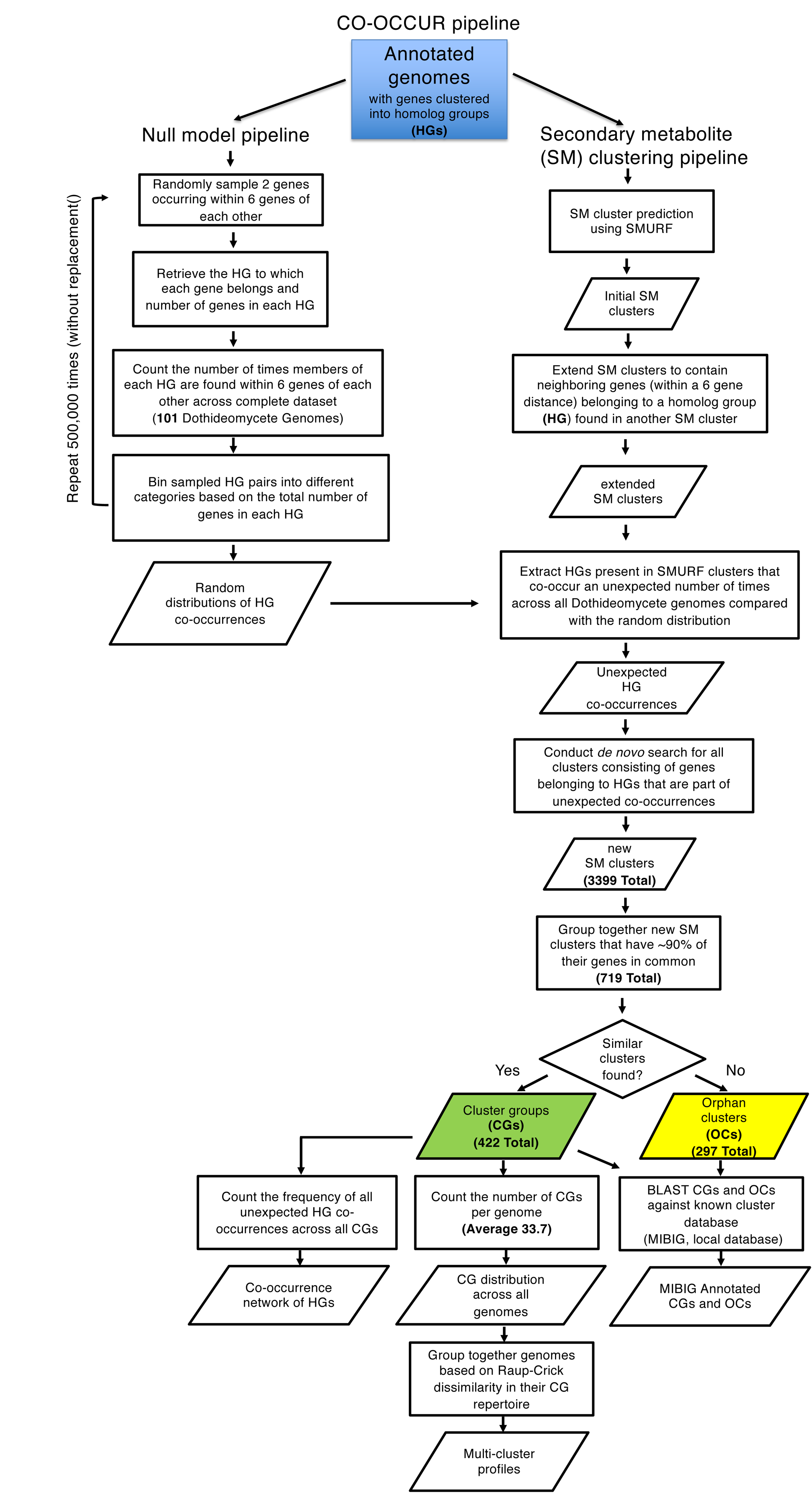

**Figure SB.**
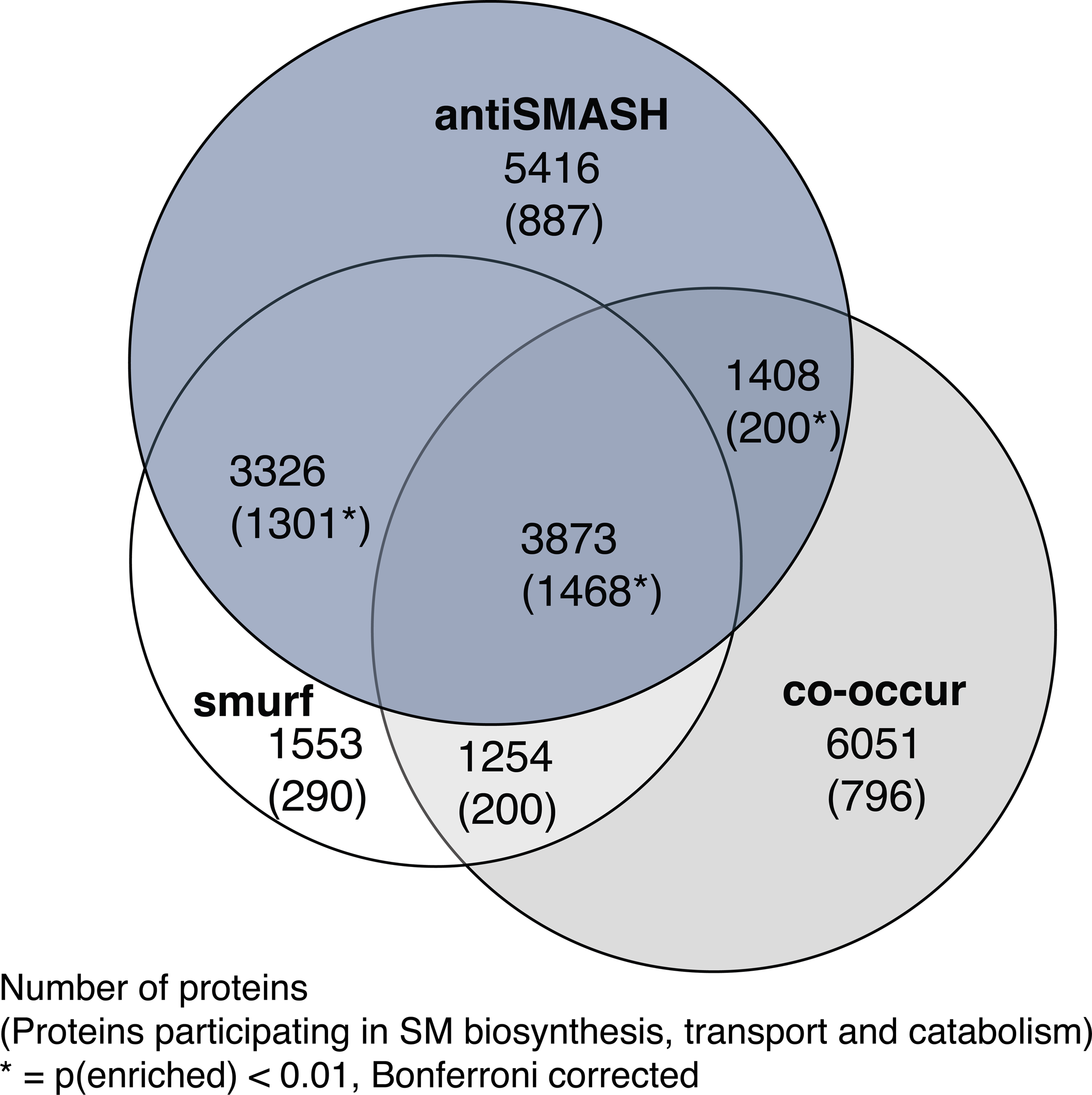

**Figure SC.**
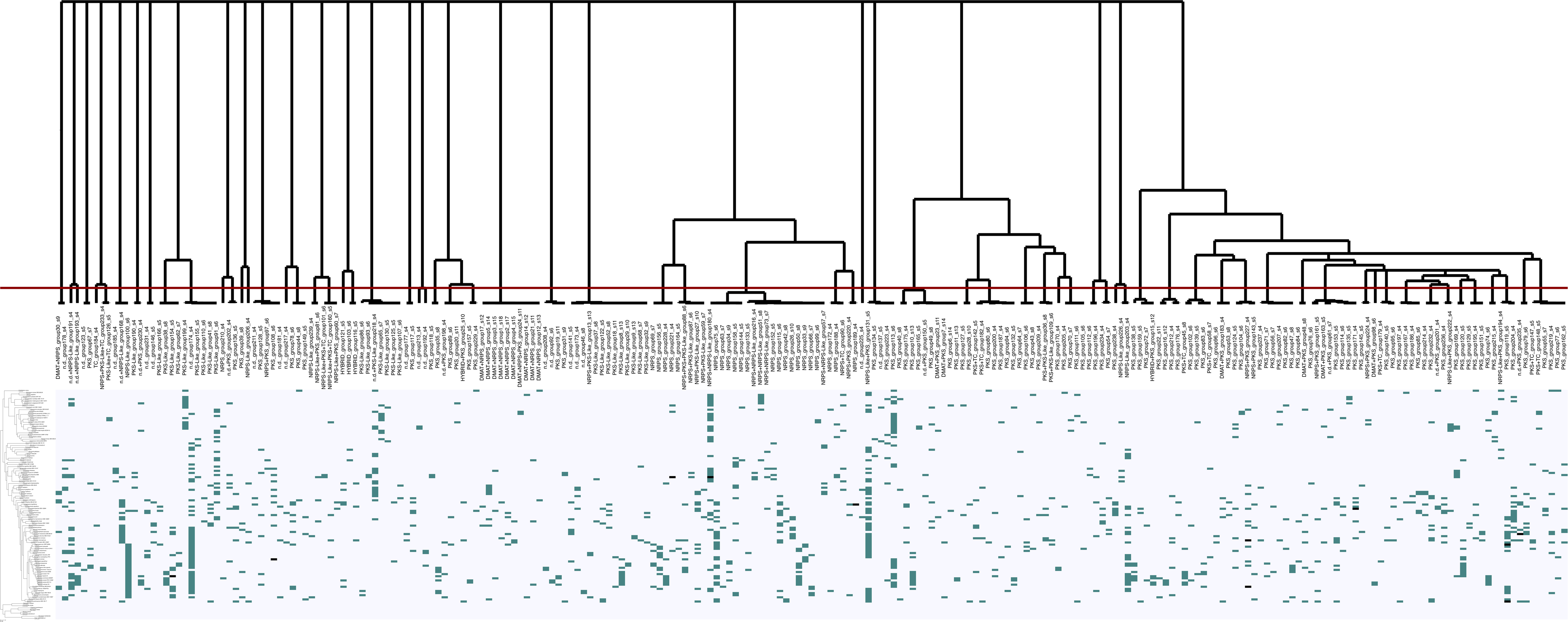

**Figure SD.**
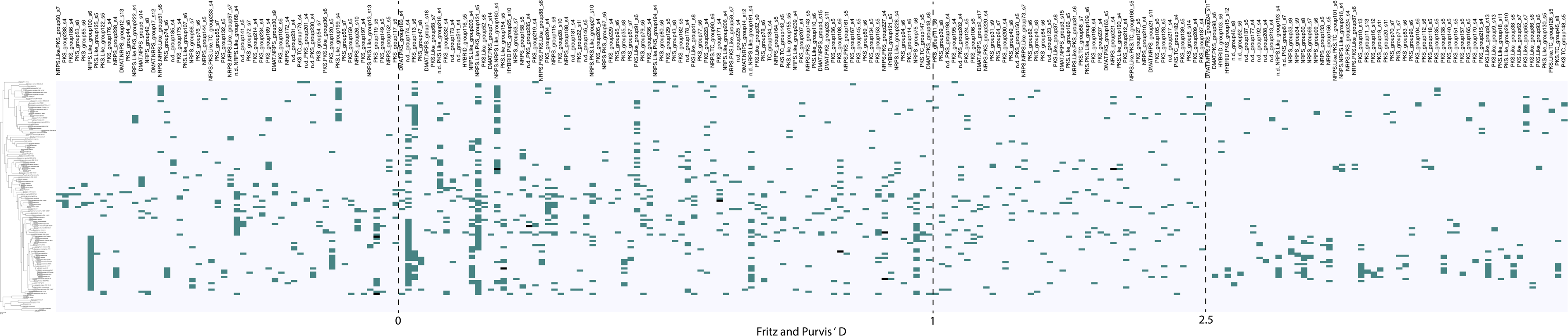

**Figure SE.**
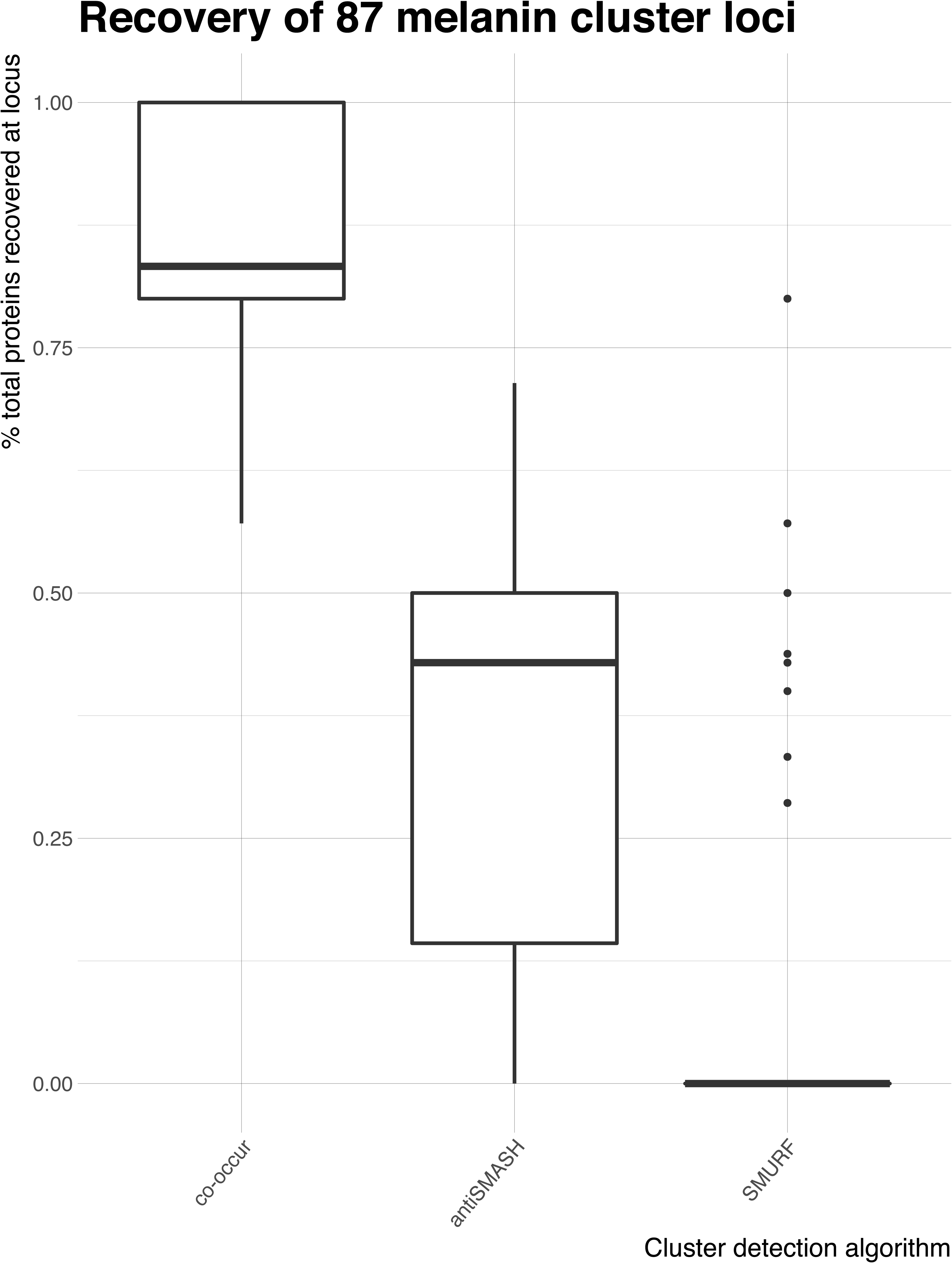

**Figure SF.**
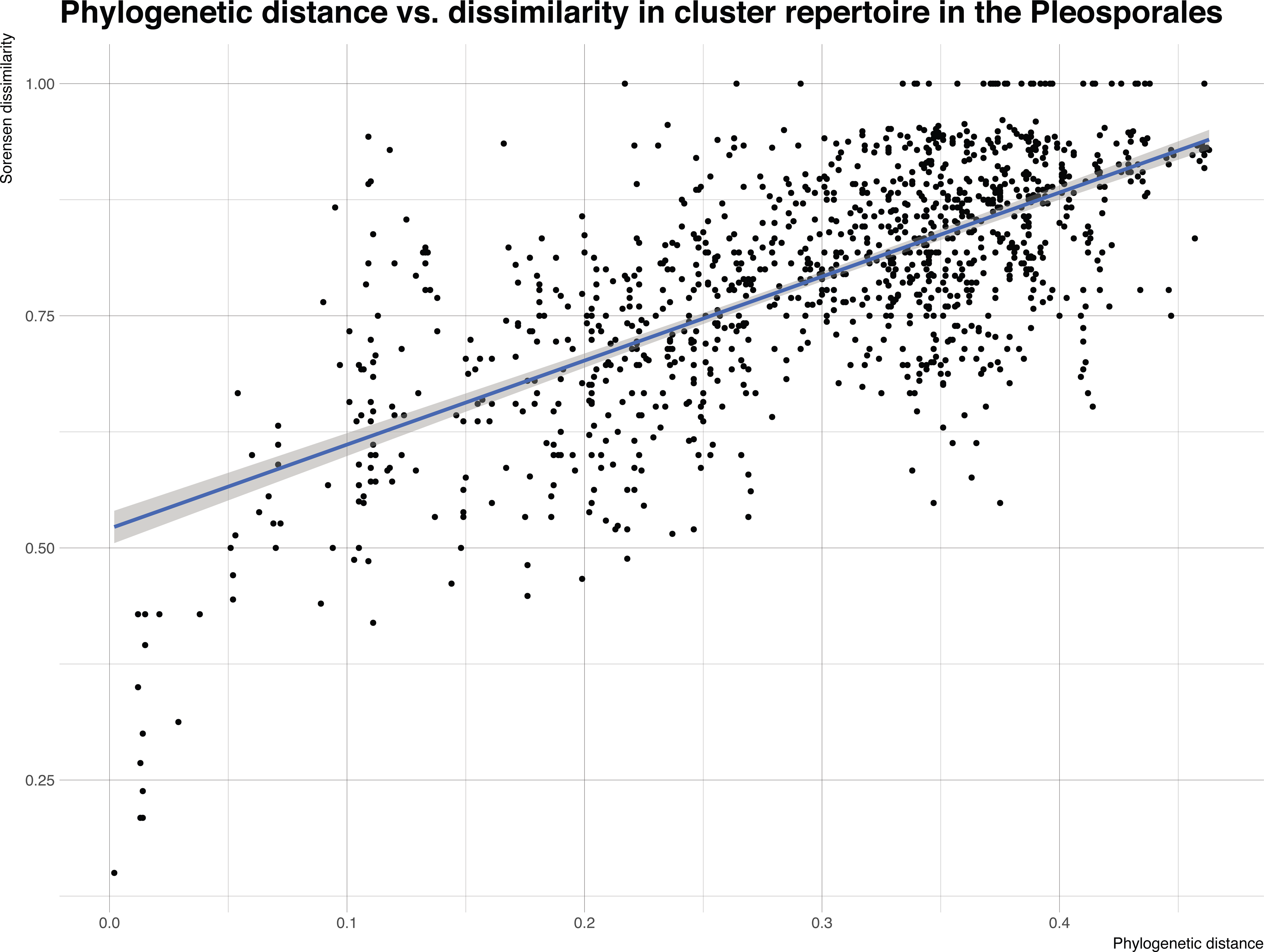

**Figure SG.**
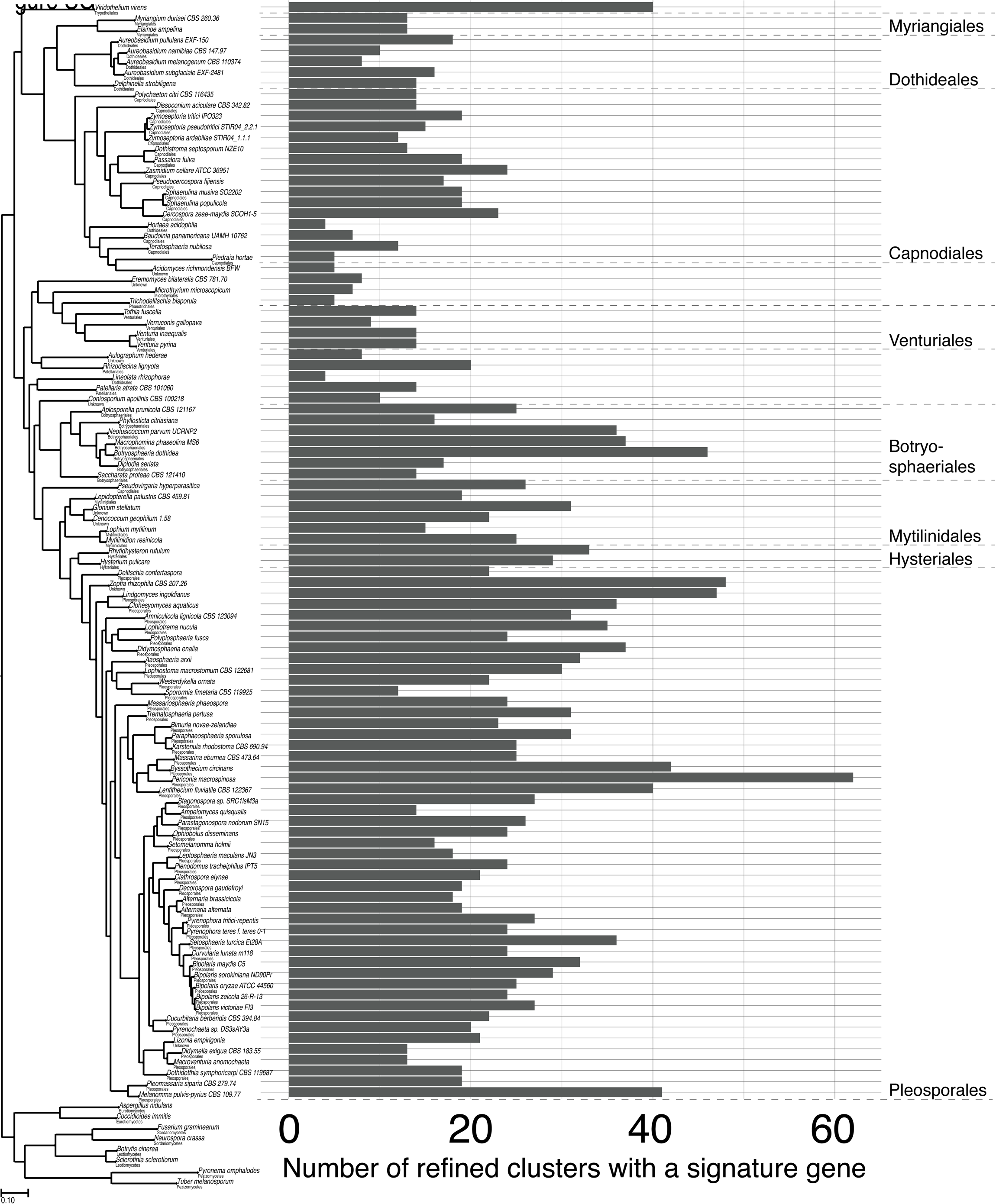

**Figure SH.**
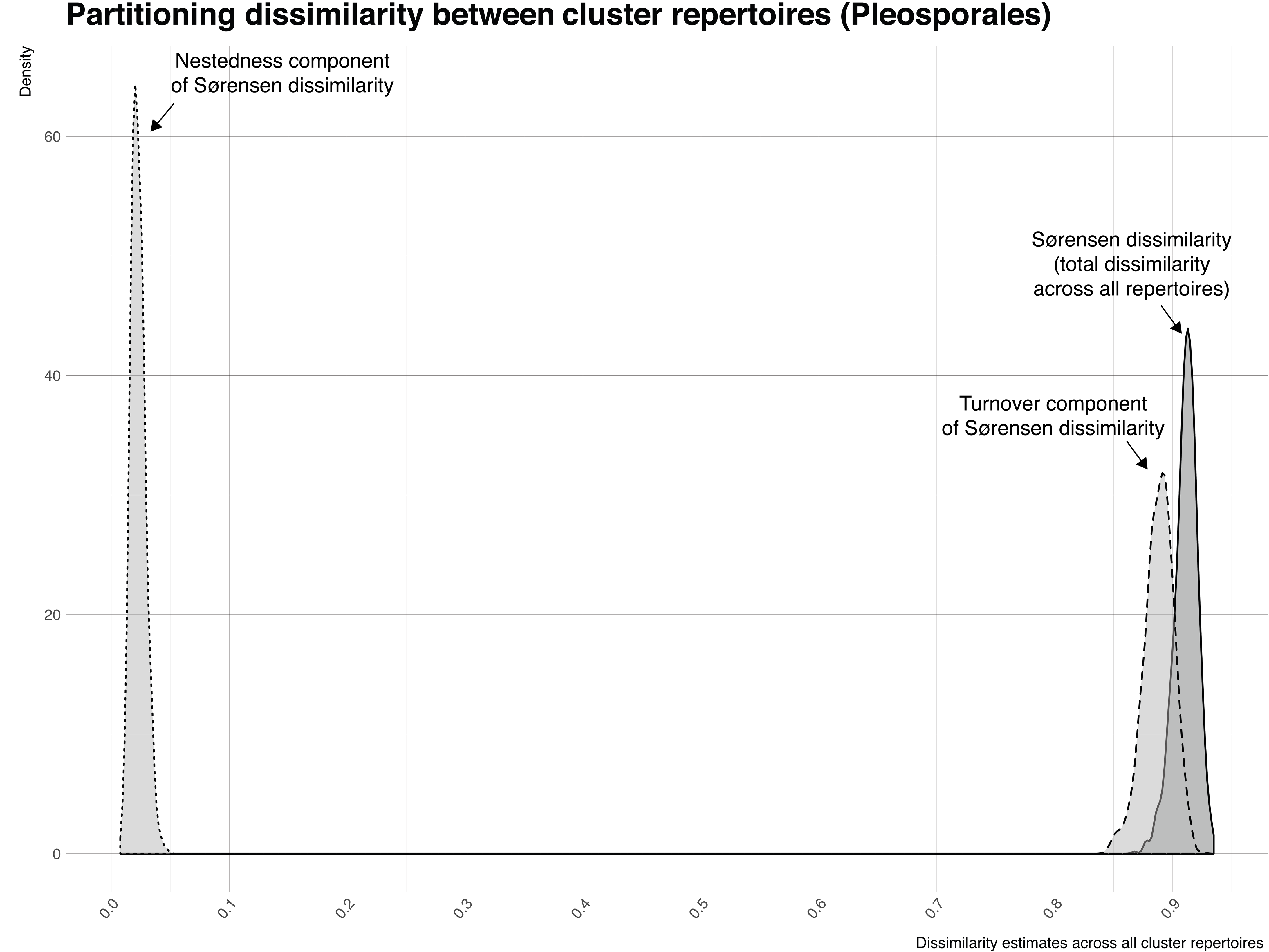

**Figure SI.**
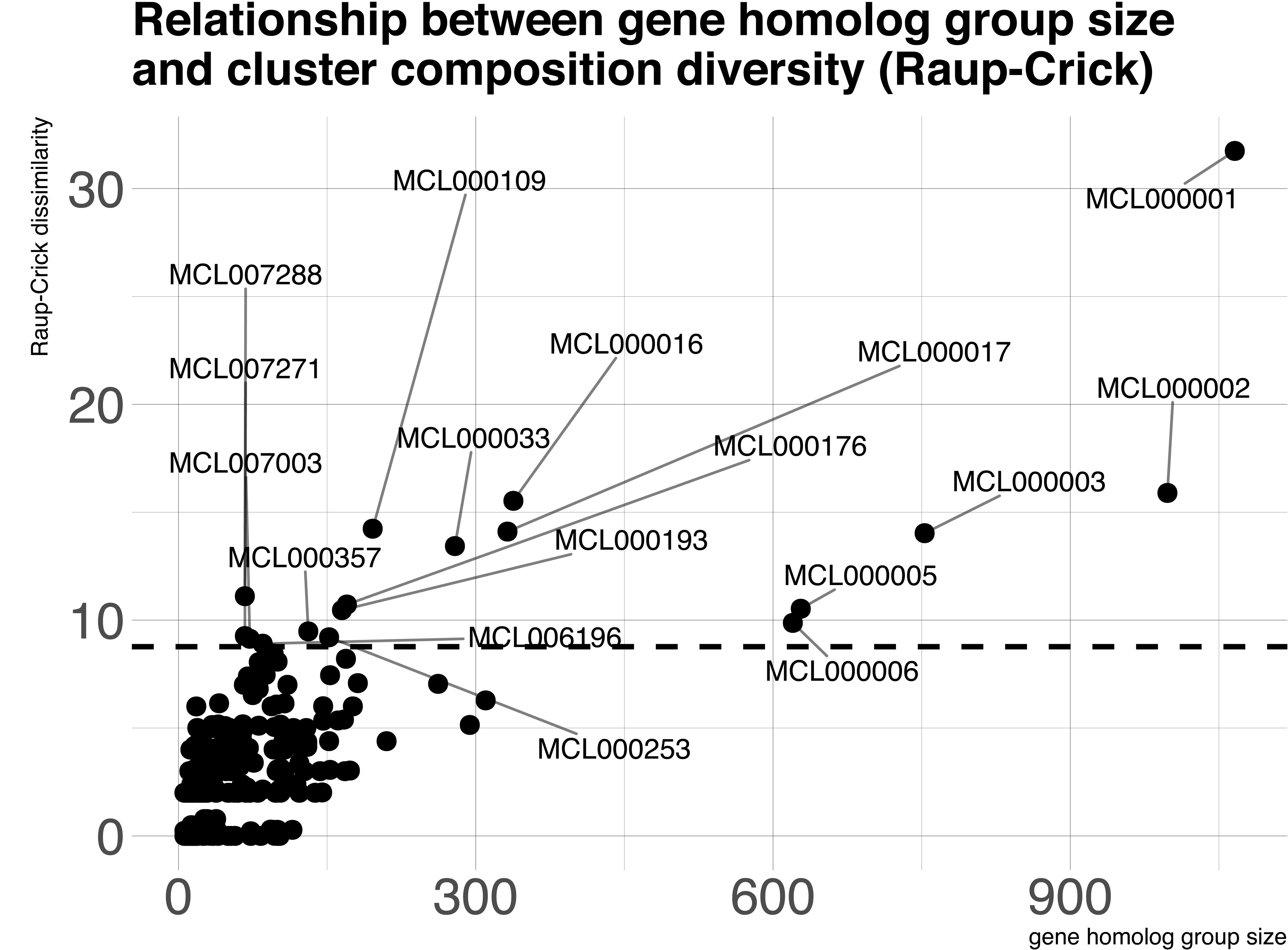

## References

1. Agrawal AA, Hastings AP, Johnson MT, Maron JL, Salminen JP. Insect herbivores drive real-time ecological and evolutionary change in plant populations. Science. 2012;338:6103.

2. Akimitsu K, Tsuge T, Kodama M, Yamamoto M, Otani H. Alternaria host-selective toxins: determinant factors of plant disease. Journal of General Plant Pathology. 2014;80:2.

3. Ali A, Caggia S, Matesic DF, Khan SA. Chaetoglobosin K, an Akt pathway inhibitor, prevents proliferation and migration of prostate carcinoma cells. 2015.

4. Baselga A, Orme CD L. betapart: an R package for the study of beta diversity. Methods in ecology and evolution. 2012;3:5.

5. Baselga, A. The relationship between species replacement, dissimilarity derived from nestedness, and nestedness. Global Ecology and Biogeography. 2012;21:12.

6. Beimforde C, Feldberg K, Nylinder S, Rikkinen J, Tuovila H, Dörfelt H, M. Gube DJ Jackson, Reitner J, Seyfullah LJ. Estimating the Phanerozoic history of the Ascomycota lineages: combining fossil and molecular data. Molecular phylogenetics and evolution. 2014;78.

7. Blin K, Wolf T, Chevrette MG, Lu X, Schwalen CJ, Kautsar SA, … Medema MH. antiSMASH 4. 0—improvements in chemistry prediction and gene cluster boundary identification. Nucleic acids research. 2017;45:W1.

8. Bok JW, Chung D, Balajee SA, Marr KA, Andes D, Nielsen KF, … Keller NP. GliZ, a transcriptional regulator of gliotoxin biosynthesis, contributes to Aspergillus fumigatus virulence. Infection and immunity. 2006;74:12.

9. Bradshaw RE, Slot JC, Moore GG, Chettri P, de Wit PJ, Ehrlich KC, … Cox MP. Fragmentation of an aflatoxin-like gene cluster in a forest pathogen. New Phytologist. 2013;198:2.

10. Chen H, Lee MH, Daub ME, Chung KR. Molecular analysis of the cercosporin biosynthetic gene cluster in Cercospora nicotianae. Molecular microbiology. 2007;64:3.

11. Choudoir MJ, Pepe-Ranney C, Buckley DH. Diversification of secondary metabolite biosynthetic gene clusters coincides with lineage divergence in Streptomyces. Antibiotics. 2018;7:1.

12. Cimermancic P, Medema MH, Claesen J, Kurita K, Brown LC W, Mavrommatis K, … Birren BW. Insights into secondary metabolism from a global analysis of prokaryotic biosynthetic gene clusters. Cell. 2014;158:2.

13. Ciuffetti LM, Manning VA, Pandelova I, Betts MF, Martinez JP. Host-selective toxins, Ptr ToxA and Ptr ToxB, as necrotrophic effectors in the Pyrenophora tritici-repentis–wheat interaction. New Phytologist. 2010 Sep;187(4):911–9.

14. Coleman JJ, Mylonakis, E. Efflux in fungi: la piece de resistance. PLoS Pathogens. 2009;5:6.

15. Condon BJ, Elliott C, González JB, Yun SH, Akagi Y, Wiesner-Hanks T, Kodama M, Turgeon BG. Clues to an evolutionary mystery: the genes for T-Toxin, enabler of the devastating 1970 Southern corn leaf blight epidemic, are present in ancestral species, suggesting an ancient origin. Molecular plant-microbe interactions. 2018 Nov 12;31(11):1154–65.

16. Crameri R, Garbani M, Rhyner C, Huitema, C. Fungi: the neglected allergenic sources. Allergy. 2014;69:2.

17. Daly, J. The host-specific toxins of Helminthosporia. Plant Infection: The Physiological and Biochemical Basis. Asada Y, Bushnell, WR, Ouchi, S., and Vance CP Berlin, Springer_verlag. 1982.

18. De Jonge R, Ebert MK, Huitt-Roehl CR, Pal P, Suttle JC, Spanner RE, … Thomma BP. Gene cluster conservation provides insight into cercosporin biosynthesis and extends production to the genus Colletotrichum. Proceedings of the National Academy of Sciences. 2018;115:24.

19. Del Carratore F, Zych K, Cummings M, Takano E, Medema MH, Breitling, R. Computational identification of co-evolving multi-gene modules in microbial biosynthetic gene clusters. Communications Biology. 2019;2:1.

20. Dhillon B, Feau N, Aerts AL, Beauseigle S, Bernier L, Copeland A, … LaButti KM. Horizontal gene transfer and gene dosage drives adaptation to wood colonization in a tree pathogen. Proceedings of the National Academy of Sciences. 2015;112:11.

21. Firn RD, Jones CG. Natural products–a simple model to explain chemical diversity. Natural product reports. 2003;20:4.

22. Fritz SA, Purvis, A. Selectivity in mammalian extinction risk and threat types: a new measure of phylogenetic signal strength in binary traits. Conservation Biology. 2010;24:4.

23. Fujii I, Yoshida N, Shimomaki S, Oikawa H, Ebizuka, Y. An iterative type I polyketide synthase PKSN catalyzes synthesis of the decaketide alternapyrone with regio-specific octa-methylation. Chemistry biology. 2005;12:12.

24. Gardiner DM, Cozijnsen AJ, Wilson LM, Pedras MS, Howlett BJ. The sirodesmin biosynthetic gene cluster of the plant pathogenic fungus Leptosphaeria maculans. Molecular microbiology. 2004 Sep;53(5):1307–18.

25. Gardiner DM, Waring P, Howlett BJ. The epipolythiodioxopiperazine (ETP) class of fungal toxins: distribution, mode of action, functions and biosynthesis. Microbiology. 2005;151:4.

26. Glassmire AE, Jeffrey CS, Forister ML, Parchman TL, Nice CC, Jahner JP, … Leonard MD. Intraspecific phytochemical variation shapes community and population structure for specialist caterpillars. New Phytologist. 2016;212:1.

27. Gluck-Thaler E, Slot JC. Specialized plant biochemistry drives gene clustering in fungi. The ISME journal. 2018;12:7.

28. Gluck-Thaler E, Vijayakumar V, Slot JC. Fungal adaptation to plant defences through convergent assembly of metabolic modules. Molecular ecology. 2018;27:24.

29. Goodwin SB, M’Barek SB, Dhillon B, Wittenberg AH, Crane CF, Hane JK, … Antoniw, J. Finished genome of the fungal wheat pathogen Mycosphaerella graminicola reveals dispensome structure, chromosome plasticity, and stealth pathogenesis. PLoS genetics. 2011;7:6.

30. Grandaubert J, Lowe RG, Soyer JL, Schoch CL, Van de Wouw AP, Fudal I, … Linglin, J. Transposable element-assisted evolution and adaptation to host plant within the Leptosphaeria maculans-Leptosphaeria biglobosa species complex of fungal pathogens. BMC genomics. 2014;15:1.

31. Grigoriev IV, Nikitin R, Haridas S, Kuo A, Ohm R, Otillar R, Riley R, Salamov A, Zhao X, Korzeniewski F, Smirnova T. MycoCosm portal: gearing up for 1000 fungal genomes. Nucleic acids research. 2014 Jan 1;42(D1):D699–704.

32. Hane JK, Rouxel T, Howlett BJ, Kema GH, Goodwin SB, Oliver RP. A novel mode of chromosomal evolution peculiar to filamentous Ascomycete fungi. Genome biology. 2011;12:5.

33. Haridas S, Albert R, Binder M, Bloem J, LaButti K, Salamov A, Andreopoulos B, Baker, SE, Barry K, Bills G, Bluhm, BH, Cannon C, Castanera R, Culley, DE, Daum C, Ezra D, González, JB, Henrissat B, Kuo A, Liang C, Lipzen A, Lutzoni F, Magnuson J, Mondo S, Nolan M, Ohm, RA, Pangilinan J, Park, H-J, Ramírez L, Alfaro M, Sun H, Tritt A, Yoshinaga Y, Zwiers L-H, Turgeon BG, Goodwin SB, Spatafora JW, Crous PW, Grigoriev IV. 101 Dothideomycetes genomes: a test case for predicting lifestyles and emergence of pathogens. Studies in Mycology, in press.

34. Hartmann FE, McDonald BA, Croll, D. Genome-wide evidence for divergent selection between populations of a major agricultural pathogen. Molecular ecology. 2018;27:12.

35. Hoffmeister D, Keller NP. Natural products of filamentous fungi: enzymes, genes, and their regulation. Natural product reports. 2007;24:2.

36. Holeski LM, Hillstrom ML, Whitham TG, Lindroth RL. Relative importance of genetic, ontogenetic, induction, and seasonal variation in producing a multivariate defense phenotype in a foundation tree species. Oecologia. 2012;170:3.

37. Holeski LM, Keefover-Ring K, Bowers MD, Harnenz ZT, Lindroth RL. Patterns of phytochemical variation in Mimulus guttatus (yellow monkeyflower). Journal of chemical ecology. 2013;39:4.

38. Horn BW. Ecology and population biology of aflatoxigenic fungi in soil. Journal of Toxicology-Toxin Reviews 2003;22:2–3.

39. Hsieh TC, Ma KH, Chao, A. iNEXT: an R package for rarefaction and extrapolation of species diversity (H ill numbers). Methods in Ecology and Evolution. 2016;7:12.

40. Huerta-Cepas J, Forslund K, Coelho LP, Szklarczyk D, Jensen LJ, Von Mering C, Bork, P. Fast genome-wide functional annotation through orthology assignment by eggNOG-mapper. Molecular biology and evolution. 2017;34:8.

41. Huerta-Cepas J, Szklarczyk D, Forslund K, Cook H, Heller D, Walter MC, … Jensen LJ. eggNOG 4. 5: a hierarchical orthology framework with improved functional annotations for eukaryotic, prokaryotic and viral sequences. Nucleic acids research. 2016;44:D1.

42. Jiang C, Song J, Zhang J, Yang, Q. New production process of the antifungal chaetoglobosin A using cornstalks. Brazilian journal of microbiology. 2017;48:3.

43. Kasahara K, Miyamoto T, Fujimoto T, Oguri H, Tokiwano T, Oikawa H, … Fujii, I. Solanapyrone synthase, a possible Diels–Alderase and iterative type I polyketide synthase encoded in a biosynthetic gene cluster from Alternaria solani. ChemBioChem. 2010;11:9.

44. Kaur, S. Phytotoxicity of solanapyrones produced by the fungus Ascochyta rabiei and their possible role in blight of chickpea (Cicer arietinum). Plant Science. 1995;109:1.

45. Keller NP. Translating biosynthetic gene clusters into fungal armor and weaponry. Nature chemical biology. 2015;11:9.

46. Khaldi N, Seifuddin FT, Turner G, Haft D, Nierman WC, Wolfe KH, Fedorova ND. SMURF: genomic mapping of fungal secondary metabolite clusters. Fungal Genetics and Biology. 2010;47:9.

47. Kurmayer R, Blom JF, Deng L, Pernthaler J. Integrating phylogeny, geographic niche partitioning and secondary metabolite synthesis in bloom-forming Planktothrix. The ISME journal. 2015;9:4.

48. Kursar TA, Dexter KG, Lokvam J, Pennington RT, Richardson JE, Weber MG, … Coley PD. The evolution of antiherbivore defenses and their contribution to species coexistence in the tropical tree genus Inga. Proceedings of the National Academy of Sciences. 2009;106:43.

49. Larsson, J. Eulerr: Area-Proportional Euler and Venn Diagrams with Ellipses. 2018. R package version 3.

50. Li D, Baldwin IT, Gaquerel E. Navigating natural variation in herbivory-induced secondary metabolism in coyote tobacco populations using MS/MS structural analysis. Proceedings of the National Academy of Sciences. 2015;112:30.

51. Lind AL, Wisecaver JH, Lameiras C, Wiemann P, Palmer JM, Keller NP, … Rokas A. Drivers of genetic diversity in secondary metabolic gene clusters within a fungal species. PLoS biology. 2017;15:11.

52. Manning VA, Pandelova I, Dhillon B, Wilhelm LJ, Goodwin SB, Berlin AM, … Holman WH. Comparative genomics of a plant-pathogenic fungus, Pyrenophora tritici-repentis, reveals transduplication and the impact of repeat elements on pathogenicity and population divergence. G3: Genes, Genomes, Genetics. 2013;3:1.

53. Menke J, Dong Y, Kistler HC. Fusarium graminearum Tri12p influences virulence to wheat and trichothecene accumulation. Molecular plant-microbe interactions. 2012;25:11.

54. Mizushina Y, Kamisuki S, Kasai N, Shimazaki N, Takemura M, Asahara H, … Sugawara F. A plant phytotoxin, solanapyrone A, is an inhibitor of DNA polymerase β and λ. Journal of Biological Chemistry. 2002;277:1.

55. Newman AG, Townsend CA. Molecular characterization of the cercosporin biosynthetic pathway in the fungal plant pathogen Cercospora nicotianae. Journal of the American Chemical Society. 2016;138:12.

56. Nielsen JC, Grijseels S, Prigent S, Ji B, Dainat J, Nielsen KF, … Nielsen J. Global analysis of biosynthetic gene clusters reveals vast potential of secondary metabolite production in Penicillium species. Nature microbiology. 2017;2:6.

57. Ohm RA, Feau N, Henrissat B, Schoch CL, Horwitz BA, Barry KW, Condon BJ, Copeland AC, Dhillon B, Glaser F, Hesse CN. Diverse lifestyles and strategies of plant pathogenesis encoded in the genomes of eighteen Dothideomycetes fungi. PLoS Pathogens. 2012 Dec;8(12).

58. Oksanen J, Kindt R, Legendre P, O’Hara B, Stevens MH, Oksanen MJ, Solymos P, Wagner H. The vegan package. Community ecology package. 2007;10.

59. Olarte RA, Menke J, Zhang Y, Sullivan S, Slot JC, Huang Y, … Bushley KE. Chromosome rearrangements shape the diversification of secondary metabolism in the cyclosporin producing fungus Tolypocladium inflatum. BMC genomics. 2019;20:1.

60. Oliver RP, Friesen TL, Faris JD, Solomon PS. Stagonospora nodorum: from pathology to genomics and host resistance. Annual review of phytopathology. 2012;50.

61. Orme D, Freckleton R, Thomas G, Petzoldt T, Fritz S, Isaac N, … Pearse W. Caper: comparative analyses of phylogenetics and evolution in R. R package version 0.5. 2012.

62. Pandelova I, Figueroa M, Wilhelm LJ, Manning VA, Mankaney AN, Mockler TC, Ciuffetti LM. Host-selective toxins of Pyrenophora tritici-repentis induce common responses associated with host susceptibility. PLoS One. 2012;7:7.

63. Patron NJ, Waller RF, Cozijnsen AJ, Straney DC, Gardiner DM, Nierman WC, Howlett BJ. Origin and distribution of epipolythiodioxopiperazine (ETP) gene clusters in filamentous ascomycetes. BMC Evolutionary Biology. 2007;7:1.

64. Proctor RH, McCormick SP, Kim HS, Cardoza RE, Stanley AM, Lindo L, … Alexander NJ. Evolution of structural diversity of trichothecenes, a family of toxins produced by plant pathogenic and entomopathogenic fungi. PLoS pathogens. 2018;14:4.

65. Reynolds HT, Vijayakumar V, Gluck-Thaler E, Korotkin HB, Matheny PB, Slot JC. Horizontal gene cluster transfer increased hallucinogenic mushroom diversity. Evolution letters. 2018;2:2.

66. Ruibal C, Gueidan C, Selbmann L, Gorbushina AA, Crous PW, Groenewald JZ, … Staley JT. Phylogeny of rock-inhabiting fungi related to Dothideomycetes. Studies in Mycology. 2009;64.

67. Schoch CL, Crous PW, Groenewald JZ, Boehm EW A, Burgess TI, De Gruyter J, … Harada, Y. A class-wide phylogenetic assessment of Dothideomycetes. Studies in mycology. 2009;64.

68. Schümann J, Hertweck C. Molecular basis of cytochalasan biosynthesis in fungi: gene cluster analysis and evidence for the involvement of a PKS-NRPS hybrid synthase by RNA silencing. Journal of the American Chemical Society. 2007;129:31.

69. Shannon P, Markiel A, Ozier O, Baliga NS, Wang JT, Ramage D, … Ideker T. Cytoscape: a software environment for integrated models of biomolecular interaction networks. Genome research. 2003;13:11.

70. Shi-Kunne X, Faino L, van den Berg GC, Thomma BP, Seidl MF. Evolution within the fungal genus Verticillium is characterized by chromosomal rearrangement and gene loss. Environmental microbiology. 2018;20:4.

71. Slot JC. Fungal gene cluster diversity and evolution. In Advances in genetics (Vol. 100, pp. 141–178). Academic Press. 2017.

72. Slot JC, Gluck-Thaler E. Metabolic gene clusters, fungal diversity, and the generation of accessory functions. Current opinion in genetics development. 2019;58.

73. Song W, Qiao X, Chen K, Wang Y, Ji S, Feng J, … Ye M. Biosynthesis-based quantitative analysis of 151 secondary metabolites of licorice to differentiate medicinal Glycyrrhiza species and their hybrids. Analytical chemistry. 2017;89:5.

74. Spatafora JW, Bushley KE. Phylogenomics and evolution of secondary metabolism in plant-associated fungi. Current opinion in plant biology. 2015;26.

75. Spatafora JW, Owensby CA, Douhan GW, Boehm EW, Schoch CL. Phylogenetic placement of the ectomycorrhizal genus Cenococcum in Gloniaceae (Dothideomycetes). Mycologia. 2012;104:3.

76. Suetrong S, Boonyuen N, Pang KL, Ueapattanakit J, Klaysuban A, Sri-indrasutdhi V, … Jones EG. A taxonomic revision and phylogenetic reconstruction of the Jahnulales (Dothideomycetes), and the new family Manglicolaceae. Fungal Diversity. 2011;51:1.

77. Theobald S, Vesth TC, Rendsvig JK, Nielsen KF, Riley R, de Abreu LM, … Hoof JB. Uncovering secondary metabolite evolution and biosynthesis using gene cluster networks and genetic dereplication. Scientific reports. 2018;8:1.

78. Turgeon BG, Baker SE. Genetic and genomic dissection of the Cochliobolus heterostrophus Tox1 locus controlling biosynthesis of the polyketide virulence factor T-toxin. Advances in genetics. 2007;57.

79. Vesth TC, Nybo JL, Theobald S, Frisvad JC, Larsen TO, Nielsen KF, … Gladden JM. Investigation of inter-and intraspecies variation through genome sequencing of Aspergillus section Nigri. Nature Genetics. 2018;50:12.

80. Villani A, Proctor RH, Kim HS, Brown DW, Logrieco AF, Amatulli MT, … Susca A. Variation in secondary metabolite production potential in the Fusarium incarnatum-equiseti species complex revealed by comparative analysis of 13 genomes. BMC genomics. 2019;20:1.

81. Walton JD. Host-selective toxins: Agents of compatibility. Plant Cell. 1996;8:10.

82. Walton JD, Panaccione DG. Host-selective toxins and disease specificity: perspectives and progress. Annual review of phytopathology. 1993;31:1.

83. Wang G, Liu Z, Lin R, Li E, Mao Z, Ling J, … Xie B. Biosynthesis of antibiotic leucinostatins in bio-control fungus Purpureocillium lilacinum and their inhibition on Phytophthora revealed by genome mining. PLoS pathogens. 2016;12:7.

84. Wang JS, Tang, L. Epidemiology of aflatoxin exposure and human liver cancer. Journal of Toxicology: Toxin Reviews. 2004;23:2–3.

85. Weinhold A, Ullah C, Dressel S, Schoettner M, Gase K, Gaquerel E, … Baldwin IT. O-acyl sugars protect a wild tobacco from both native fungal pathogens and a specialist herbivore. Plant physiology. 2017;174:1.

86. Wight WD, Kim KH, Lawrence CB, Walton JD. Biosynthesis and role in virulence of the histone deacetylase inhibitor depudecin from Alternaria brassicicola. Molecular plant-microbe interactions. 2009;22:10.

87. Wijayawardene NN, Hyde KD, Lumbsch HT, Liu JK, Maharachchikumbura SS, Ekanayaka AH, … Phookamsak R. Outline of ascomycota: 2017. Fungal Diversity. 2018;88:1.

88. Wisecaver JH, Slot JC, Rokas, A. The evolution of fungal metabolic pathways. PLoS Genetics. 2014;10:12.

89. Wolpert TJ, Dunkle LD, Ciuffetti LM. Host-selective toxins and avirulence determinants: what’s in a name?. Annual review of phytopathology. 2002;40:1.

90. Xu YJ, Luo F, Li B, Shang Y, Wang C. Metabolic conservation and diversification of Metarhizium species correlate with fungal host-specificity. Frontiers in microbiology. 2016;7.

91. Ziemert N, Lechner A, Wietz M, Millán-Aguiñaga N, Chavarria KL, Jensen PR. Diversity and evolution of secondary metabolism in the marine actinomycete genus Salinispora. Proceedings of the National Academy of Sciences. 2014;111:12.

92. Züst T, Heichinger C, Grossniklaus U, Harrington R, Kliebenstein DJ, Turnbull LA. Natural enemies drive geographic variation in plant defenses. Science. 2012;338:6103.

